# CHROMOMETHYLTRANSFERASE3/ KRYPTONITE maintain the *sulfurea* paramutation in *Solanum lycopersicum*

**DOI:** 10.1101/2021.07.01.450764

**Authors:** Cláudia Martinho, Zhengming Wang, Andrea Ghigi, Sarah Buddle, Felix Barbour, Antonia Yarur, Quentin Gouil, Sebastian Müller, Maike Stam, Chang Liu, David C. Baulcombe

## Abstract

Paramutation involves the transfer of a repressive epigenetic mark from a silent allele to an active homologue and, consequently, non-Mendelian inheritance. In tomato the *sulfurea* (*sulf*) paramutation is associated with a high level of CHG hypermethylation in a region overlapping the transcription start site of the *SlTAB2* gene that affects chlorophyll synthesis. The CCG sub-context hypermethylation is under-represented at this region relative to CTG or CAG implicating the CHROMOMETHYLTRANSFERASE3 (CMT3) in paramutation at this locus. Consistent with this interpretation, loss of *CMT*3 function leads to loss of the *sulf* chlorosis, the associated CHG hypermethylation and paramutation. Loss of *KRYPTONITE (KYP)* histone methyl transferase function has a similar effect linked to reduced H3K9me2 at the promoter region of *SlTAB2* and a shift in higher order chromatin structure at this locus. Mutation of the largest subunit of RNA polymerase V (*PolV*) in contrast does not affect *sulf* paramutation. These findings indicate the involvement of a CMT3/KYP dependent feedback rather than the PolV-dependent pathway leading to RNA directed DNA methylation (RdDM) in the maintenance of paramutation.

## Introduction

Paramutation causes non-Mendelian inheritance in which an epigenetic mark at a silenced *paramutagenic* allele transfers to and silences an active *paramutable* homologue (1–3). In plants the DNA is methylated at the paramutated (silenced) sequence region, but this modification is not sufficient to mediate paramutation, because many methylated loci do not show paramutation. Early examples of paramutation in maize (1), pea (4) and tomato (5) have striking phenotypes based on pigmentation or gross morphology. However, from genome-wide DNA methylation analyses, there is now evidence that paramutation-like effects may be more widespread (6–8). In some examples the trigger for paramutation may be hybridisation between distantly related varieties or even species (6–8).

Like all heritable epigenetic mutations, paramutation involves separate mechanisms to establish a molecular mark at the paramutable locus and to mediate its maintenance through cycles of cell division and sexual reproduction. Additionally, a defining characteristic of paramutation is the interaction between the participating alleles that, in many models, involves the RdDM pathway (1, 2).

RdDM is not, however, exclusive to paramutation. At many genomic loci it mediates DNA methylation in transposable elements (TEs) and repetitive regions (9). It involves RNA polymerase IV (PolIV) transcription of the target DNA and RNA-dependent RNA polymerase (RDR2) conversion of the transcript into double stranded RNA. DCL3 trims the double stranded RNA into 24 nucleotide (nt) fragments and, after unwinding, the single stranded 24nt siRNAs are loaded into Argonaute protein 4 (AGO4). This AGO4 nucleoprotein is then guided by Watson-Crick base pairing to a RNA polymerase V (PolV)-generated scaffold RNA and it recruits DNA methyltransferases, including DRM2, that catalyse methylation of adjacent DNA cytosines (9).

The RdDM factors identified in paramutation screens in maize include the RDR2 orthologue (Mediator of Paramutation 1 (MOP1)) which is required to maintain and establish paramutation at multiple maize loci, including the classic paramutation example in the *booster1* (b1) locus (10). Maize genetic screens have also uncoveremultiple PolIV subunits including MOP2/RMR7 (RDP2), MOP3, RMR6, RMR7, RDP1 and RDP2a as affecting paramutation (1). RPD2a has the potential to integrate PolV complexes (11) Consistent with involvement of the RdDM pathway, there may be abundant 24nt siRNAs and high DNA methylation at the target loci. These 24nt siRNAs could, in principle, diffuse within the nucleus and mediate the allelic interaction in paramutation.

However, the picture from genetic screens is incomplete. There is no clear evidence for involvement of the PolV subunits, AGO proteins, DNA methyltransferases or other factors in the downstream part of the RdDM pathway or the 21nt siRNAs or AGO6 associated with the initial establishment phase (12). Some maize studies also found it difficult to reconcile these findings with an siRNA-based model for paramutation (13). Paramutation at maize *purple plant1* (*pl1*), for example, can be established in mutants which lack siRNAs, including, RMR1 a Snf2 protein which affects the stability of nascent transcripts (14). Conversely, mutations in the RMR12 chromodomain helicase DNA-binding 3 (CHD3) protein orthologous to Arabidopsis (*Arabidopsis thaliana*) PICKLE release the *pl1* maize paramutation but in a manner that probably promotes the incorporation of nucleosomes without affecting sRNA accumulation (15). These various findings and observations do not necessarily rule out the involvement of siRNAs or RdDM in paramutation but they do point to the gaps in current models. Therefore, we are investigating *sulf* paramutation in tomato (16). Our reasoning is that *sulf* paramutation may be mechanistically similar to the maize examples but the genetic architecture of different plants would provide a new perspective on the factors involved.

The *sulf-*mediated leaf chlorosis phenotype is due to hypermethylation in a Differentially Methylated Region1 (DMR1) at the 5’ region of *SlTAB2 (TAB2)*, a gene which affects the synthesis of chlorophyll in tomato plants (16). *TAB2* is silenced between 0-100% in the F1 progeny of *sulf* × wild type (WT) plants (17). In crosses between variegated *sulf* lines and their parental WT lines paramutation is poorly penetrant in F1 (<12%) (17). In F2, the percentage of variegation among heterozygous also varies, further showing that *sulf* has incomplete penetrance (5, 17). These differences in paramutation penetrance have been linked with the existence of different sulfurea epi-alleles with different degrees of paramutagenicity (5, 17).

Allele-specific DNA methylation analysis at DMR1 in interspecific *sulf × S. pimpinellifolium* F1 hybrids, revealed high methylation levels in the CHG context in the *de nov*o paramutated allele (16). In tomato, CHG-methylation is maintained by CHROMOMETHYLTRANSFERASE3a (*SlCMT3a*, hereafter referred as CMT3) (18). Here we show that *sulf* paramutation requires the self-reinforcing loop involving CHG DNA methylation by the DNA methyl transferase CMT3 and H3K9 dimethylation by the histone methyltransferase KRYPTONITE (KYP) at the silenced locus which correlate with changes in chromosome compartment. Mutation of the largest RNA polymerase V subunit (NRPE1) does not affect *sulf*. Our findings suggest that *sulf* paramutation in tomato might be independent of the canonical RdDM pathway and that CHG methylation is required.

## Results

### CMT3 maintains the *sulf* paramutation

The *sulf* phenotype results from *TAB2* hypermethylation in a DMR1 close to the TSS (19). The severity of leaf chlorosis is directly proportional to the levels of DMR1 DNA methylation and thus far is not associated with variation in DNA sequences (16). Here we classify *sulf* epialleles based on their correlated phenotype and epigenotype (Table 1) (16). Despite displaying high differential DNA methylation at the CG context (∼50%), DMR1 is unusual in that it has a very high level of CHG hypermethylation (∼60%) (16) consistent with the involvement of CMT3 (18). To test this hypothesis, we assayed methylation at the different CHG subcontexts in DMR1 (Fig. 1A) (19) in leaf tissue excised from plants bearing *TAB2*^*+*^ (WT cv. M82) or *TAB2*^*sulf*^ (*sulf*). CMT3 preferentially methylates CAG and CTG motifs over CCG in multiple plant species, including tomato (18, 20). In *TAB2*^*sulf*^ the percentage of CCG methylation in DMR1 was lower than at CAG and CTG (Fig. 1B), consistent with the involvement of *CMT3* in *TAB2*^*sulf*^. The subcontext bias was less pronounced in the gene body of *TAB2* than in DMR1 (Fig. 1B). It was also less pronounced in random non-DMR regions of similar size within the pericentric heterochromatin of chromosome 2 (Fig. 1B, Table S1). To further test this possibility, we took advantage of the plants bearing CRISPR-mediated deletion in the *SlCMT3*a gene - *cmt3*a (referred to as *cmt3*) (18). The heterozygous *CMT3*/*cmt3* was crossed to *CMT3*/*CMT3* plants with the *TAB2*^*sulf*^ epiallele (Fig. 1C). In the F1 generation, we selected plants heterozygous for *cmt3* and, from McrBC-qPCR (16), for the *TAB2*^*sulf/+*^ epialleles characteristic of the epiheterozygous state at DMR1 (Fig. 1C, S1A). The epiheterozygous state is characteristic of a *sulf* allele with low paramutation penetrance.

**Table 1.** Epiallele classification employed in this study.

| Epigenotype | Genotype | DNA methylation / Phenotype | Paramutated | Identification |
| --- | --- | --- | --- | --- |
| ***TAB2*<sup>+</sup>** | M82 DNA sequence | <50% methylation, Green | No | phenotyping McrBC-qPCR |
| ***TAB2*<sup>sulf/+</sup>** | M82 DNA sequence | 50% methylation, Green (e.g. F1s) | No | phenotyping McrBC-qPCR |
| ***TAB2*<sup>sulf</sup>** | M82 DNA sequence | >50% methylation, Variegated leaf chlorosis | Yes | phenotyping McrBC-qPCR |

**Figure 1.**
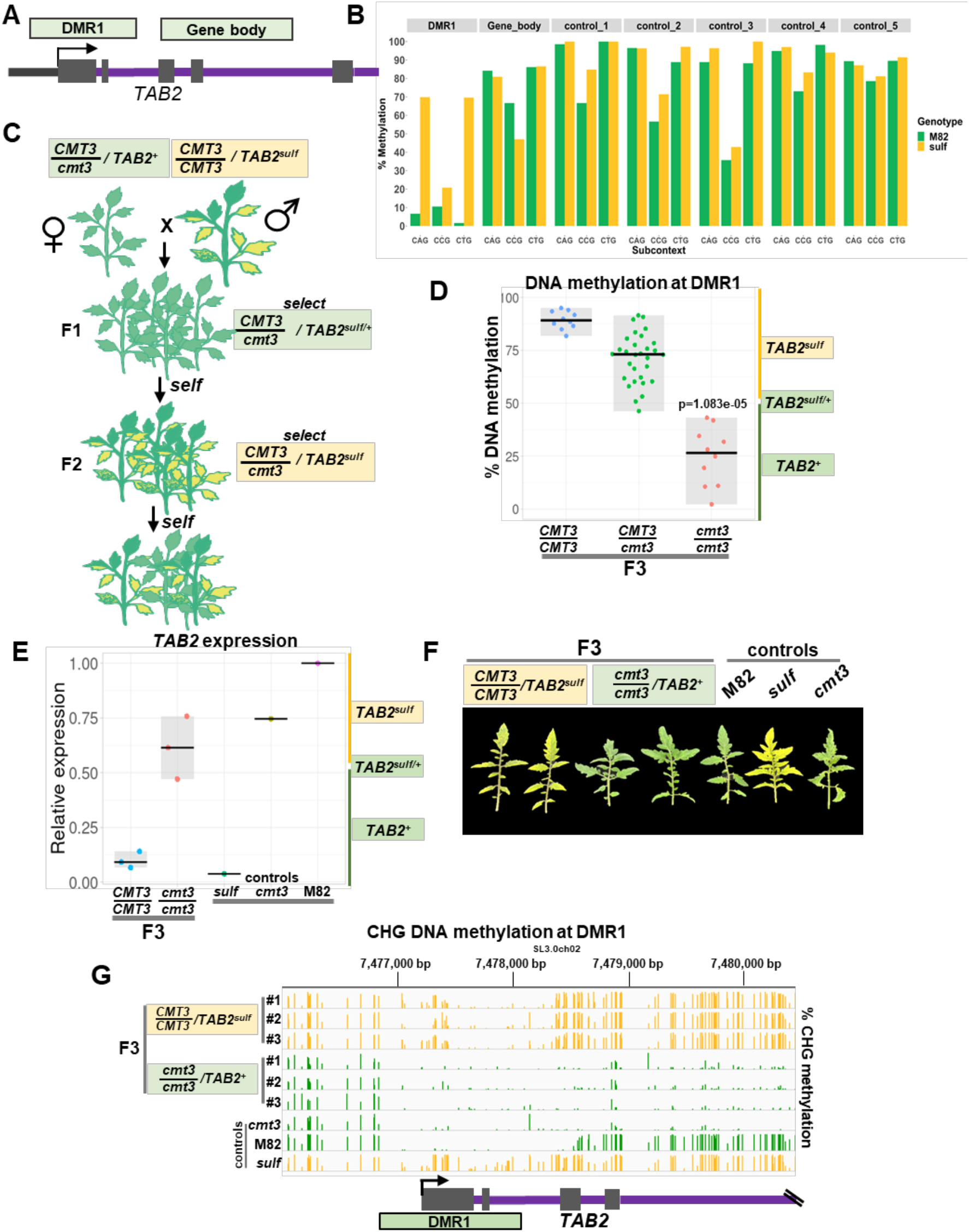
CMT3 maintains *sulf*. **A–** Diagram illustrates *TAB2* locus and relative DMR1 and gene body position. These regions were used for analysis in Fig. 1B – refer to Table S1 for precise coordinates. **B–** Mean % DNA methylation in leaf tissue at CHG subcontext in DMR1, *TAB2* gene body and five random control regions within SL3.0ch02. “M82”-*S. lycopersicum* cv. M82 *TAB2*^*+*^ (green) n=2; and “*sulf”* - *S. lycopersicum cv. Lukullus TAB2*^*sulf*^ (yellow) n=2. Coordinates are listed in Table S1. **C–** Diagram illustrates crossing scheme used to generate F3 populations (F3 pedigree). **D–** Jittered dots depict % of DNA Methylation at DMR1 in individual plants determined by McrBC-qPCR. Plants denote F3 siblings – 4-week-old leaf tissue. The summary of the data is shown as horizontal line indicating the median. Grey boxes illustrate the data range. × axis refers to *CMT3* genotypes. *CMT3/CMT3 TAB2*^*sulf*^ n=10; *CMT3/cmt3 TAB2*^*sulf*^ n=30; *cmt3/cmt3 TAB2*^*+*^ n=10. p-value *cmt3/cmt3* versus *CMT3/CMT3* was calculated employing a Mann-Whitney-Wilcoxon test. **E–** Jittered dots depict relative *TAB2* expression in individual plants (4-week-old leaf tissue) normalised to the geometric mean of the expression of two reference genes (Table S3). The summary of the data is shown as horizontal line indicating the median. Grey boxes illustrate the data range. F3 plants: *CMT3/ CMT3 TAB2*^*sulf*^ n=3; *cmt3/cmt3 TAB2*^*sulf*^ n=3. Controls: M82-(*S. lycopersicum cv. M82) CMT3/ CMT3 TAB2*^*+*^ *n=1; sulf -* (*S. lycopersicum cv. Lukullus) CMT3/CMT3 TAB2*^*sulf*^ *n=1; cmt3 – cmt3/ cmt3 TAB2*^*+*^ n=1. **F–** 2 month-old leaves. F3 plants and controls: same as in Fig. 1E. F3 plants and controls: same as in Fig. 1E. **G–** *TAB2* IGV screenshot of bisulphite sequencing data exhibiting CHG DNA methylation (4-week-old leaf tissue), range [0-100]; F3 plants and controls: same as in Fig. 1E. Green tracks refer to green leaf phenotype and yellow tracks refer to plants which display chlorosis. **A** and **C-G–**Yellow boxes refer to plants displaying *sulf* chlorosis and green boxes refer to green plants.

In the F2 generation, only *CMT3* homozygotes or heterozygotes displayed the *TAB2*^*sulf*^ phenotype and epigenotype (>50% methylation by McrBC-qPCR) (Fig. S1B). Thus, we conclude that DMR1 hypermethylation and CMT3 are required for establishment or maintenance of *sulf* paramutation. A similar genetic dependence on *CMT3* for the *TAB2*^*sulf*^ epiallele was observed in the F3 progeny of *CMT3*/*cmt3* F2 plants *TAB2*^*sulf*^ (Fig 1D). As in the F2, the homozygous *cmt3* F3 plants were green and had lower DMR1 DNA methylation levels than their *CMT3* siblings (Fig 1D) indicative of the epiallele *TAB2*^*+*^.

Further support for a link between *sulf* chlorosis, DNA methylation, and CMT3 level comes from the analysis of DMR1 DNA methylation in the heterozygous *CMT3*/*cmt3* F2 and F3 plants. The distribution of DMR1 DNA methylation in these plants was skewed towards lower values than in *CMT3* homozygotes (Fig S1B, Fig 1D) and, in the F2, a smaller proportion of plants were *TAB2*^*sulf*^ and chlorosis in *CMT3/cmt3* (∼17%) than in *CMT3*/*CMT3* (∼24%) (Fig S1B). Many of the green F2 *CMT3/cmt3* heterozygous plants may have inherited *TAB2*^*sulf*^ and *TAB2*^*sulf/+*^ but with incomplete maintenance of *TAB2*^*sulf*^ and reversion to *TAB2*^*+*^ due to reduced dosage of *CMT3*.

Consistent with the low DNA methylation levels of DMR1 in the F3 *cmt3* backgrounds, *TAB2* expression in these plants was higher than in the F3 *CMT3* siblings (Fig. 1E) although not as high as in the *CMT3* or *cmt3* lines with *TAB2*^*+*^ that had not been crossed with *sulf* (Fig. 1E). The homozygous *cmt3* plants lacked *sulf* chlorosis whereas their F3 *CMT3* siblings were chlorotic (Fig. 1F, S2).

More detailed analysis of the F3 DNA methylation pattern by bisulphite sequencing analysis confirmed that, compared to M82, the CHG DNA methylation in *sulf* plants bearing the *TAB2*^*sulf*^ allele was higher in DMR1extending from 212 bp in the 5’ upstream region into intron 2 at 852 bp downstream of the TSS (19) (Fig 1G). In the *cmt3 TAB2*^*+*^ F3 progeny the hyper CHG DMR1 was lost (Fig. 1G). There was also CHG hypomethylation of the transcribed DNA but no indication that this gene body DNA methylation influenced paramutation (Fig. 1G). From these patterns we conclude that *sulf* is associated with CMT3-mediated CHG hypermethylation of DMR1.

In the CG context the *sulf* hyper DMR was restricted to the DMR1 region but the association with *sulf* was weaker than with CHG (Fig. S3). The degree of hyper CG methylation was markedly lower in the *cmt3 TAB2*^*+*^ F3 progeny than in the *CMT3 TAB2*^*sulf*^ siblings (Fig S3). The CHH context did not exhibit significant hypermethylation in the TAB2 DNA of *sulf* plants (Fig. S3). There was a region of CHH hypomethylation on the upstream side of DMR1 (Fig. S3) corresponding to a Differentially Methylated Region2 (DMR2) (16) but it was not correlated with the *sulf* phenotype in the F3 progeny: the hypomethylation was lost to a varying extent irrespective of whether the plants were *CMT3* and *TAB2*^*sulf*^ or *cmt3* and *TAB2*^*+*^ (Fig. S3). These results indicate that CHG methylation patterns, but not CG or CHH, primarily govern the *sulf* phenotype.

The *cmt3* mutants lose DMR1 hypermethylation and *sulf* chlorosis but, in principle, they could retain other epigenetic marks with the potential to re-establish *TAB2*^*sulf*^ in a *CMT3* background. To assess this possibility, we backcrossed a F2 *cmt3*/*cmt3 TAB2*^*+*^ plant with one of the highest level of methylation (27%) (Fig. S3B) with a M82 *TAB2*^*+*^ allele (Fig. S4A) and quantified DNA methylation at DMR1 using McrBC-qPCR in the BC1 and two generations of selfed progeny (Fig. S4A). None of these plants had DMR1 DNA methylation levels higher than 40% (Fig. S4B-D) and plants were not chlorotic (Fig. S4E). Bisulphite sequencing confirmed lower levels of CHG methylation at DMR1 in backcrossed plants than *sulf* controls (Fig. S5) suggesting that *sulf* memory is therefore *CMT3*- dependent and RdDM is not sufficient to maintain *sulf*.

### NRPE1 and *sulf* paramutation

To assess a possible role of an RdDM cofactor we crossed *sulf* plants with a mutant in an RdDM component DNA-directed RNA polymerase V subunit 1 (NRPE1) (Fig. 2A) (20) and then selfed F1 plants that were epiheterozygous (*TAB2*^*sulf/+*^) (Fig. 2A, S6A) for two generations. The mutant has a deletion of two codons in the *NRPE1* coding sequence, reduced 24nt siRNAs and hypomethylation at RdDM loci (20). In the F2 and F3 progeny the DMR1 DNA methylation profiles were similar in all *NRPE1* genotypes (Fig. 2B, S6B). In the F2 the mean level was close to 50% with a distribution from 0-100% (Fig. S6B). In the F3, all of the plants had the *TAB2*^*sulf*^ epiallele since DMR1 methylation was above 50% in all individuals regardless of their *NRPE1* genotype (Fig. 2B). On average, the F3 *nrpe1* plants had 20% lower DMR1 DNA methylation levels than their *NRPE1* F3 siblings (Fig. 2B) but this reduction did not suppress silencing of *TAB2* expression (Fig. 2C) or the *sulf* chlorosis (Fig. 2D, S7). From these data we conclude that reduced NRPE1 function has a minor effect on DMR1 DNA methylation but that PolV is not an essential cofactor of *sulf* silencing.

**Figure 2.**
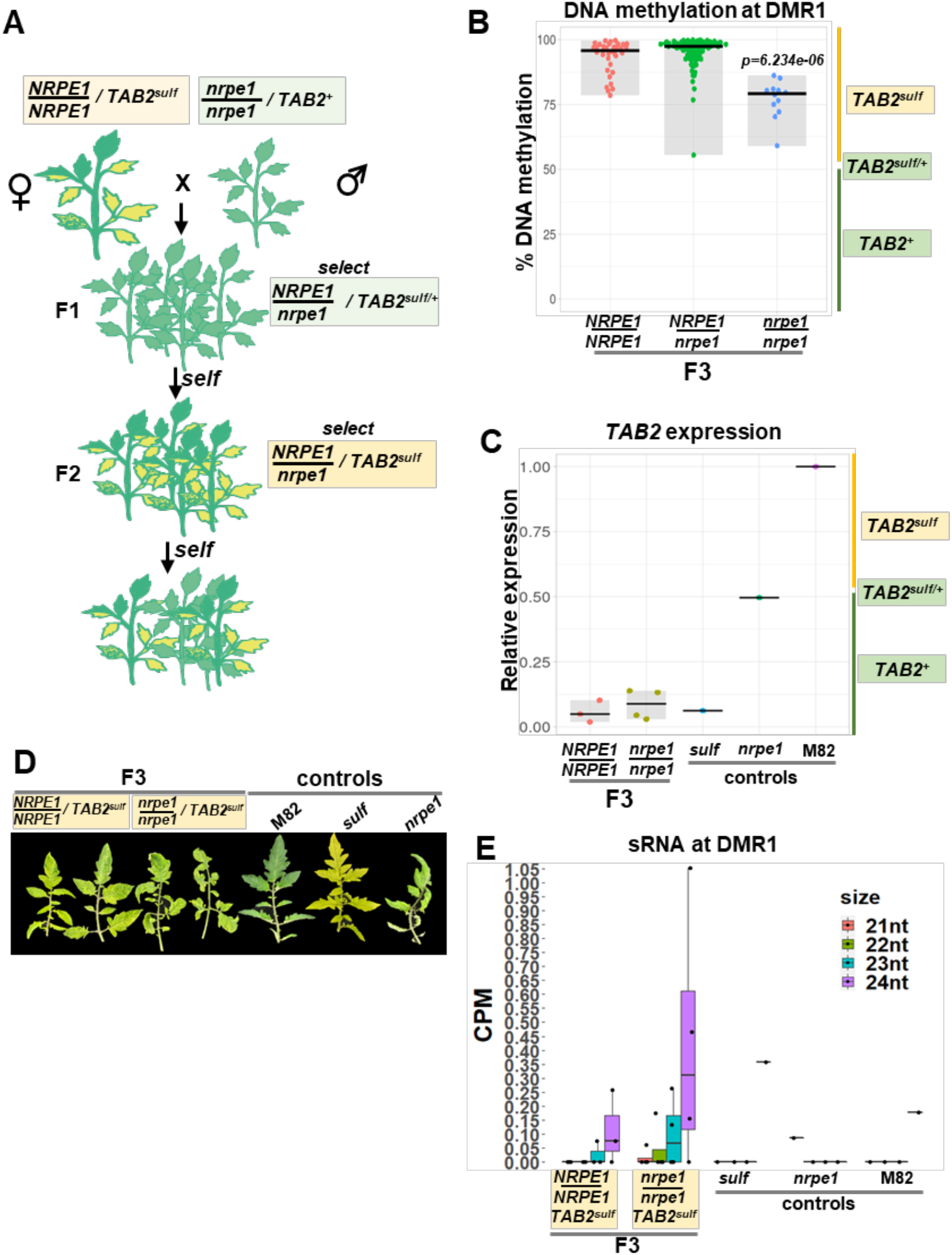
sulf maintenance remains unaffected in nrpe1 mutants. **A–** Diagram illustrates crossing scheme used to obtain F3 populations (F3 pedigree). **B–** Jittered dots depict individual plants’ % DNA Methylation at DMR1 determined by McrBC-qPCR. Plants denote F3 siblings – 4- week-old leaf tissue. The summary of the data is shown as horizontal line indicating the median. Grey boxes illustrate the data range. × axis refers to *NRPE1* genotypes. *NRPE1/NRPE1 TAB2*^*sulf*^ n=35; *NRPE1/nrpe1 TAB2*^*sulf*^ n=94; *nrpe1/nrpe1 TAB2*^*sulf*^ n=12. p-value *nrpe1/ nrpe1* versus *NRPE1/NRPE1* was calculated employing a Mann-Whitney-Wilcoxon test. **C–** Jittered dots depict relative *TAB2* expression in individual plants (4-week-old leaf tissue) normalised to the geometric mean of the expression of two reference genes (Table S3). The summary of the data is shown as horizontal line indicating the median. Grey boxes illustrate the data range. F3 plants: *NRPE1/NRPE1 TAB2*^*sulf*^ n=3; *nrpe1/nrpe1 TAB2*^*sulf*^ n=4. Controls: M82-(*S. lycopersicum cv. M82) NRPE1/NRPE1 TAB2*^*+*^ *n=1; sulf -* (*S. lycopersicum cv. Lukullus) NRPE1/NRPE1 TAB2*^*sulf*^ *n=1; nrpe1 – nrpe1/nrpe1 TAB2*^*+*^ n=1. **D–** 2 month-old leaves. F3 plants and controls: same as Fig. 2C. **E–** DMR1 siRNA size distribution in counts per million (CPM) in F3 plants and controls. (4-week-old leaf tissue). Genotypes are the same as in Fig. 2C. Jittered dots representindividual plants. The summary of the data is shown as horizontal line indicating the median. Errors bars represent the standard deviation. Genotypes are the same as in Fig. 2C. **A-E**-Yellow boxes refer to plants displaying *sulf* chlorosis and green boxes refer to green plants.

24nt siRNAs constitute a hallmark of RdDM activity (9). Given the non-essential role of NRPE1 in maintaining *sulf*, we reassessed siRNAs at the *TAB2* locus employing sRNA-sequencing. On average 24nt siRNAs accumulate more in *TAB2*^*sulf*^ than in *TAB2*^*+*^ at DMR1 (16) but at a low and variable level (Fig. S8A). Gene body sRNAs showed a more consistent upregulation in *sulf* plants (Fig. S8B).

Whole genome sRNA-sequencing data revealed that 13% of all sRNA loci were affected by NRPE1 (20) but, in F3 *NRPE1* and F3 *nrpe1* plants, the low level of DMR1 24nt sRNA were not significantly different (Fig 2E). Similarly, in *CMT3* and *cmt3* F2 plants of the crosses with *TAB2*^*sulf*^ the DMR1 sRNA of all 21-24 size classes were unaffected by genotype or *sulf* epigenotype (Fig. S9A and B). There was a reduction of gene body sRNAs in *cmt3* genotypes with *TAB2*^*+*^ (Fig. S9 A and C), but this effect did not correlate with *sulf* chlorosis: DNA methylation was abundant in plants with *TAB2*^*+*^ epigenotype (Fig. 1G and S3, compare controls *sulf* with M82). From these sRNA data it is unlikely, therefore, that DMR1 or gene body siRNAs play a role in *sulf* maintenance.

### KYP and H3K9me2 are required to maintain the *sulf* paramutation

In Arabidopsis, a self-reinforcing loop model involving H3K9me2, a repressive histone modification, can account for maintenance of CHG methylation by CMT3. In this model, KRYPTONYTE (KYP) family proteins bind CHG DNA methylation and methylate histone H3k9 tails. In turn, *CMT3* binds H3K9me2 and increases the existing level of CHG DNA methylation (21, 22). Given the signature of CMT3-dependent DNA methylation at *TAB2*, the CMT3/KYP reinforcement model predicts higher levels of H3K9me2 in *sulf* than WT tissues at DMR1.

To test this prediction, we carried out H3K9me2 ChIP followed by quantitative real time PCR (ChIP-qPCR) with oligonucleotide pairs A-F spanning different regions of the *TAB2* locus (Fig. 3A) in M82 and *sulf* plants bearing *TAB2*^*+*^ and *TAB2*^*sulf*^ epigenotypes, respectively. Consistent with our prediction, the H3K9me2 enrichment was higher in *TAB2*^*sulf*^ than in *TAB2*^*+*^ at region B in TAB2 promoter and the DMR1 overlapping regions C, D and E, including the *TAB2* TSS (Fig. 3B). In contrast, we detected no or smaller H3K9me2 differences between *TAB2*^*+*^ and *TAB2*^*sulf*^ at regions outside DMR1, including A and F and an unrelated Transposable Element (TE) (Fig. 3B). From these results we conclude that the link between CMT3 and *sulf* is associated with increased levels of H3K9me2 at DMR1.

**Figure 3.**
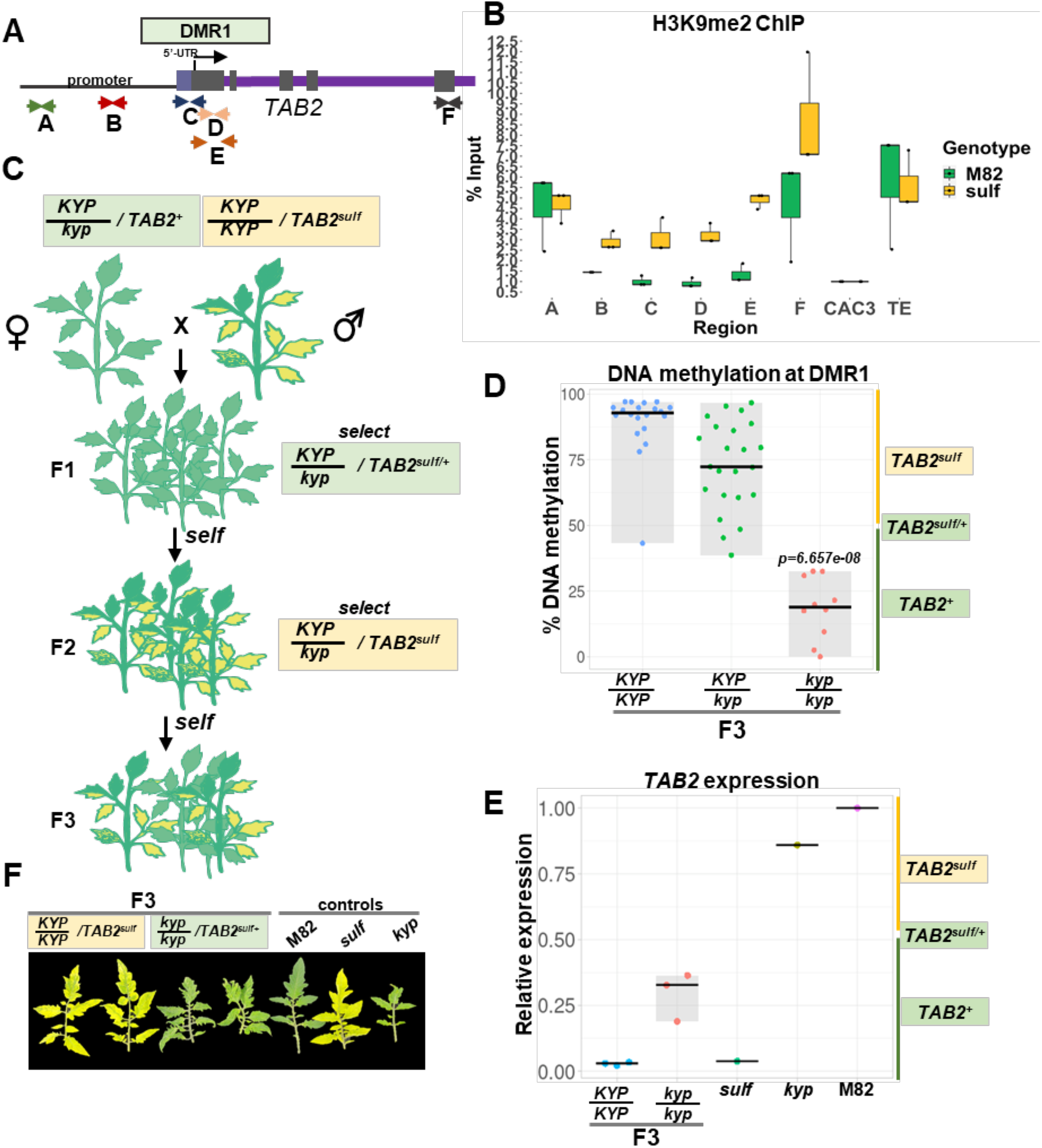
KYP maintains *sulf*. **A–** Diagram represents the relative oligonucleotide position spanning the *TAB2* locus used for ChIP-qPCR experiments in Fig. 3B. **B–** Box plot depicts H3K9me2 enrichment per % input normalised to *CAC3* reference locus determined by Chip-qPCR. Jittered dots represent different biological replicates. “M82”- *S. lycopersicum* cv. M82 TAB2^+^ (green) n= 3; “*sulf”* - *S. lycopersicum cv. Lukullus* TAB2^sulf^ (yellow) n=3. The summary of the data is shown as horizontal line indicating the median of biological replicates. Error bars represent the standard deviation. **C–** Diagram illustrates crossing scheme used to obtain F3 populations (F3 pedigree). **D–** Jittered dots depict % DNA Methylation at DMR1 individual plants determined by McrBC-qPCR. Plants denote F3 siblings. The summary of the data is shown as horizontal line indicating the median. Grey boxes illustrate the data range. × axis refers to *KYP* genotypes. F3 plants: *KYP/KYP TAB2*^*sulf*^ n=20; *KYP/kyp TAB2*^*sulf*^ n=23; *kyp/kyp TAB2*^*+*^ n=10. p-value *kyp/kyp* versus *CMT3/CMT3* was calculated employing a Mann-Whitney-Wilcoxon test. **E–** Jittered dots depict relative *TAB2* expression in individual plants normalized using the geometric mean of the expression values for two reference genes (Table S3). The summary of the data is shown as horizontal line indicating the median. Grey boxes illustrate the data range. F3 plants: *KYP/KYP TAB2*^*sulf*^ n=3; *kyp/kyp TAB2*^*+*^ n=3. Controls: M82-(*S. lycopersicum cv. M82) KYP/KYP TAB2*^*+*^ *n=1; sulf -* (*S. lycopersicum cv. Lukullus) KYP/KYP TAB2*^*sulf*^ *n=1; kyp – kyp/kyp TAB2*^*+*^ n=1. **F–** 2 month-old leaves. Genotypes are the same as in Fig. 3E. **A-E**-Yellow boxes refer to plants displaying *sulf* chlorosis and green boxes refer to green plants.

In M82 leaves bearing *TAB2*^*+*^, the H3K9me2 levels were lower close to the TSS than in more distal regions (Fig. 3B). Such low levels are exceptional in pericentromeric heterochromatin (where TAB2 resides) (18) and it is likely that these low H3K9me2 levels enable transcription of *TAB2*. Consistent with this interpretation, there were high levels of the active transcription mark H3K4me3 close to *TAB2* TSS (Fig. S10) detected by ChIP-qPCR in M82 plants at regions C, D and E. The association between histone modifications with *sulf* silencing is reinforced by reduced levels of the H3K4me3 mark in at these sites at *TAB2*^*sulf*^ (Fig. S10).

To further test the involvement of H3K9me2 we crossed *sulf* with a mutant bearing CRISPR-Cas9-mediated deletion at the gene encoding for the *S. lycopersicum* KYP (18) (Fig. 3C). The crossing strategy was parallel to that used for *cmt3* and, in the F2 and F3 progeny of the F1 *KYP/kyp* epiheterozygous *TAB2*^*sulf/+*^ (Fig. S11A), the *kyp/kyp* plants all had less than 50% DMR1 DNA methylation (Fig. S11B, Fig 3D). Accordingly, *TAB2* expression F3 *kyp* plants was higher than in the *KYP* siblings (Fig 3E), and none displayed symptoms of chlorosis (Fig 3F, S12). Only progeny that carried an active KYP allele displayed *TAB2*^*sulf*^ epigenotype, *sulf* phenotypes and low TAB2 expression (Fig 3DE), as observed in the original *TAB2*^*sulf*^ parent. The *sulf* paramutation in tomato, therefore, is associated with high levels of CHG DNA methylation and H3K9me2 at DMR1 that would be expected to influence chromatin organisation.

### Chromatin architecture changes in *sulf*

To test this hypothesis, we carried out Chromatin Conformation Capture analysis (Hi-C) in *TAB2*^*sulf*^ and *TAB2*^*+*^. Despite having distinct leaf colours, M82 and *sulf* plants showed highly similar Hi-C maps, suggesting that there are no intensive genome rearrangements or changes in chromatin contacts in *sulf* plants bearing *TAB2*^*sulf*^ epialleles (data not shown). PCA analysis of the contact matrix of chromosome 2 revealed A/B compartments that correlated to the delineation of euchromatin/heterochromatin across this chromosome (Fig. 4 A, B). This A/B compartment partition was broadly similar in the two epigenotypes and in all the samples, the genomic region containing *TAB2* belonged to the heterochromatic B compartment (Fig. 4C). However, there was a change in the PCA eigenvalue of this genomic region in all the *sulf* replicates, indicating a change in chromatin interaction between this region and the A/B compartments. By examining the interaction strength between this *TAB2*-containing region and the A compartment, we found that it was significantly weaker in the *TAB2*^*sulf*^ plants than in the *TAB2*^*+*^ plants (Fig 4D), indicating a general shift away from euchromatic regions.

**Figure 4.**
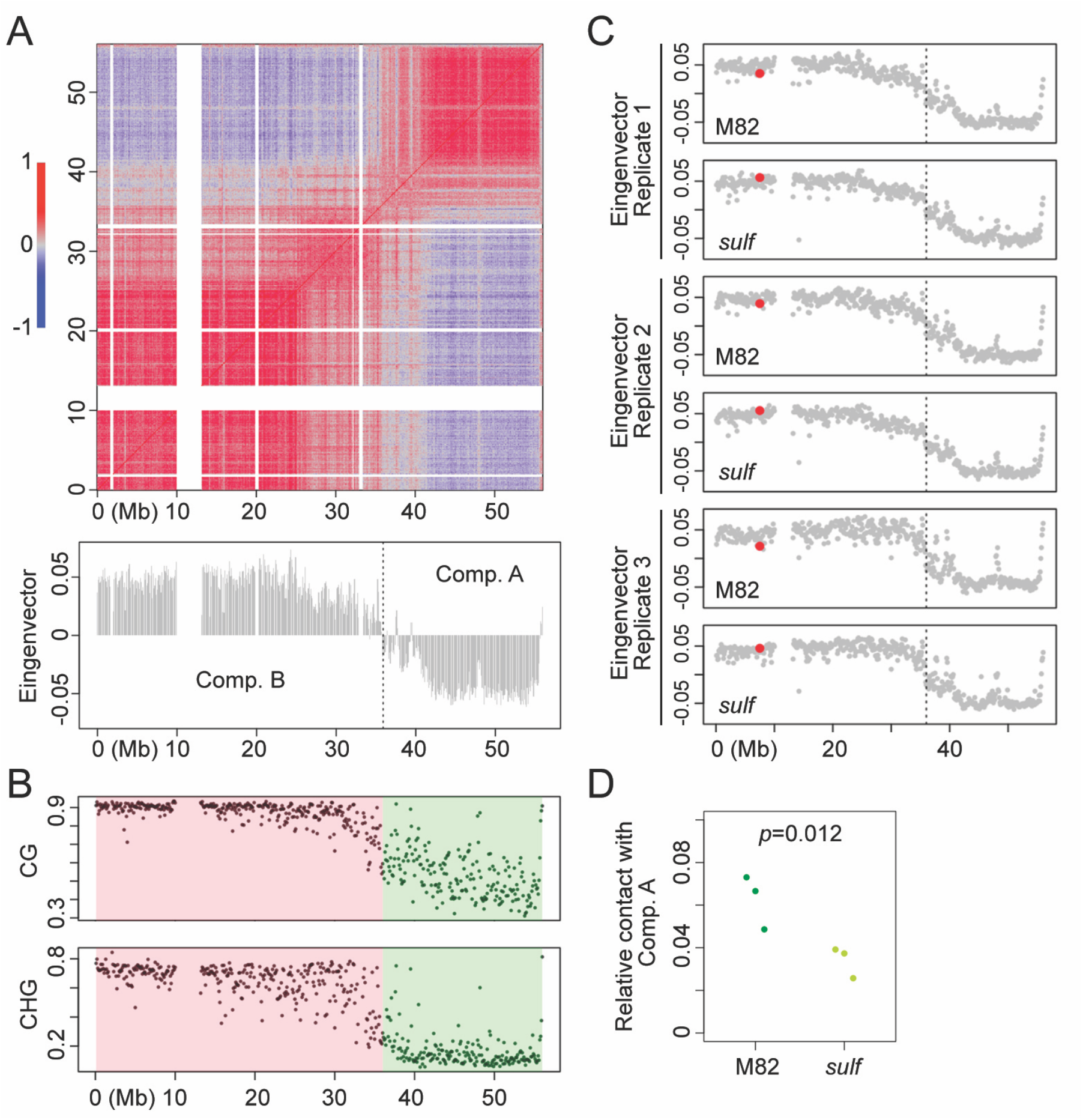
*sulf* is associated with differential genome organization. **A and B-** Annotation of A/B compartment. **A-** The Hi-C map (normalized at 100 kb resolution) of chromosome 2 from M82 (*S. lycopersicum* cv. M82 *TAB2*^*+*^) (replicate 1) is shown. The bottom plot depicts PCA (principal component analysis), from which the A/B compartment is deduced. **B-** Comparison of DNA methylation ratios of genomic regions in A (green block) and B (red block) compartments. **C-** PCA showing Hi-C maps of chromosome 2 from M82 and *sulf* (*S. lycopersicum cv. Lukullus TAB2*^*sulf*^) plants. In each plot, the red dot depicts the genomic region which contains the *TAB2* locus, the vertical line shows the A/B compartment border described in panel A. **D-** Relative chromatin contacts between the region bearing the *TAB2* locus and the A compartment. n=3. p value indicates paired t-test result.

## Discussion

We confirm here that *sulf* paramutation is associated with DMR1 that overlaps the transcriptional start site of *TAB2* (16) (Fig. 1G, S3) and we have shown additionally the involvement of CMT3/KYP (Fig. 1 and 3) and an associated change in chromatin organisation (Fig. 4). CMT3/KYP have not been previously associated with endogenous gene paramutation, although in *Arabidopsis*, a *LUC* paramutation transgene relies on multiple proteins including MET1 and CMT3/KYP (23). The involvement of CMT3 in maize is difficult to test because loss of function in the *ZMET2* and *ZMET5* homologues is not compatible with plant viability (24). However, the subcontext CHG DNA methylation pattern associated with B’, a highly paramutagenic maize allele at the b1 locus, reminisces CMT3 signatures (Fig. S13). This may suggest that *ZMET2* and *ZMET5* play a role in the maintenance of b1 paramutation in maize. High H3K9me2 levels correlate with b1 and p1 paramutagenic alleles (25, 26), consistent with the involvement of a CMT3/KYP self-reinforcing loop as at *sulf*.

The dynamics of *sulf* paramutation vary depending on the *sulf* epiallele used as parent. Some *sulf* epialleles transfer the repressive state in the F1 but, with the *sulf* epialleles used here the transfer likely occurs in the F2 (17). The *sulf* epiallele was transmitted and maintained in the *CMT3*/*CMT3* homozygous F1 or *CMT3*/*cmt3* heterozygous F1 and then progressively transferred between *TAB2* alleles in the *CMT3*/*CMT3* or *CMT3*/*cmt3* F2 but not in the *cmt3*/*cmt3* homozygous F2. While we cannot rule out that F2 plants with the *sulf* phenotype had inherited two silent alleles of *TAB2*, it is known that *TAB2*^*sulf*^ homozygotes die at the seedling stage (5) and therefore, a more likely scenario is that they germinated as *TAB2*^*sulf/+*^ and then progressed to *TAB2*^*sulf*^ through paramutation shortly after germination.

One scenario is that CMT3/KYP is required to make the *TAB2*^*+*^ chromatin sensitive to receive the paramutation silencing signal from a silenced allele (Fig. 5 – scenario 1). Alternatively, the paramutagenic allele (*sulf*) silences the paramutable allele (M82) in the *cmt3* homozygote as it would do so in the *CMT3* genotype, but the silencing is not maintained as *CMT3* is needed to either complete the *sulf* silencing process or to allow the repressed state to persist (Fig. 5 – scenario 2). In the former scenario the role of *CMT3* would be in establishment and in the latter it would be in maintenance of paramutation (Fig. 5).

**Figure 5.**
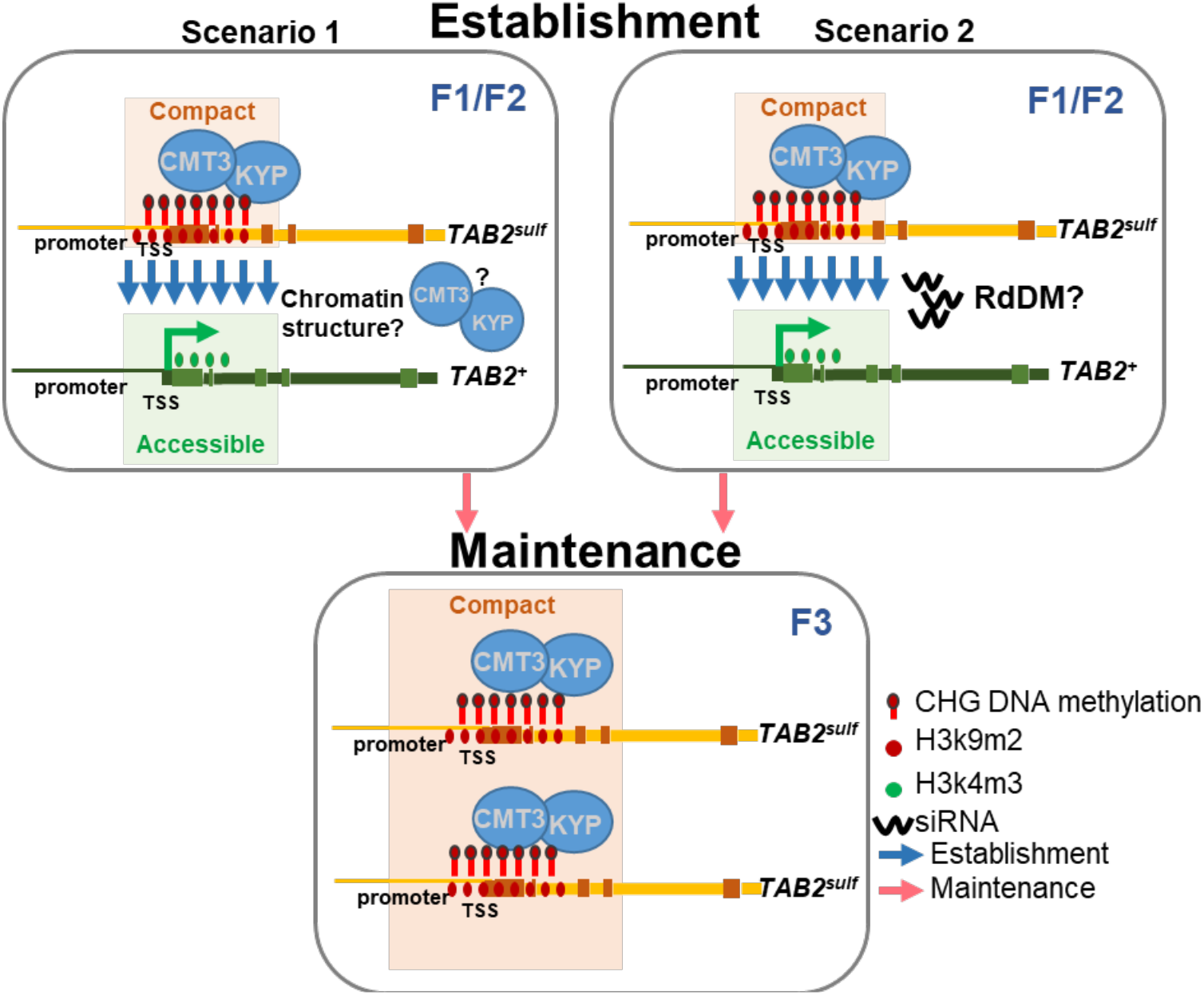
Model to explain the establishment (F1 or F2) and maintenance of the *sulf* paramutation in tomato. Scenario 1 and 2 represent alternative hypotheses to explain the establishment of the *sulf* paramutation which are not mutually exclusive. In scenario 1, the establishment of paramutation requires a CMT3/KYP-dependent compact chromatin conformation which in turn facilitates the exchange of epigenetic marks. In scenario 2, the initial round of DNA methylation at *TAB2* is initiated by RdDM and siRNAs. We show that CMT3/KYP is required either to complete silencing or/and maintain *TAB2*^*sulf*^ in subsequent cell divisions and across generations. Yellow/orange boxes represent the silenced paramutagenic epiallele *TAB2*^*sulf*^ which exists in a compact chromatin conformation. Green boxes represent the active paramutable *TAB2*^*+*^ epiallele in which chromatin exists in a less compact structure and therefore remains more accessible to the transcription machinery.

The involvement of KYP and the changes to the chromatin modifications and organisation of *TAB2* in *sulf* are consistent with all of these hypotheses. However, the finding that CMT3 is absolutely required to maintain *sulf* epigenetic memory (Figs. S4 and S5) implies a role of CMT3/KYP and the associated change in chromatin architecture in maintenance. This finding contrasts with demonstration that paramutated states persist through backcrosses with maize RdDM mutants(10). The various *CMT*3 hypotheses are not, however, mutually incompatible and its involvement in maintenance does not rule out a second role in establishment.

The accepted paradigm for establishment of paramutation invokes sRNAs in the communication between interacting alleles. In our *sulf* system, however, there is a very low level of (24nt) siRNA accumulation at *TAB2*^*sulf*^ and loss or reduced function of PolV does not lead to loss of *sulf* silencing. These findings are not easy to reconcile with simple role of RdDM in *sulf* paramutation although we cannot rule it out conclusively. It could be that there is residual PolV function in the mutant line and that *sulf* siRNA is abundant at specific developmental stages when communication between alleles takes place.

Even in maize, however, the 24nt siRNA role in paramutation also remains unclear. For instance, RMR1, and RMR7 are not required for paramutation establishment at Pl1-Rhoades despite affecting 24nt siRNA accumulation (13, 14). In addition, paramutation is affected by genetic backgrounds with normal 24nt siRNA levels (15). Furthermore, siRNA accumulation is not sufficient to direct paramutation at *b1*, indicating that other factors must be involved (27).

In summary, our results indicate that models of paramutation should accommodate the involvement of CMT3/KYP and changes in chromatin structure in at least the maintenance/memory phase if not in establishment (Fig. 5). These models should be open to the possibility that there could be locus-specific and/or species specific mechanisms of paramutation and that there could be communication between alleles by mechanisms other than through sRNA and RdDM. Further understanding of chromatin organisation and the 4D genome will also be necessary to understand how paramutation, uniquely amongst epigenetic phenomena, enables communication between alleles.

## Materials and Methods

Detailed experimental procedures are provided in SI.

## Acknowledgments

We would like to thank Melanie Steer for the invaluable horticultural support. We thank Pawel Baster, James Barlow, Neuza Duarte and Jean-Francois Popoff for technical support. We would also like to thank Christoph Lambing and Paul Fransz for the effort of troubleshooting FISH experiments which weren’t successful and could not be included in this manuscript. Finally, we would like to thank all the colleagues who supported this study through scientific discussions: Jake Harris, Hajk-Georg Drost, Sara Lopez-Gomollon and Hadi Putra. This study was supported by the BBSRC grant PDAG/387.

## Author Contributions

C.M. and D.C.B. designed the research; C.M., C.L, A.G., S.B., F.B., A.Y. and M.S. performed research; C.M., D.C.B., C.L., Z.W., Q.G., M.S. and S.M. analysed data; and C.M. and D.C.B. wrote the paper.

