## Supplemental Information for "CHROMOMETHYLTRANSFERASE3/ KRYPTONITE maintain the *sulfurea* paramutation in *Solanum lycopersicum*"

### **Supplementary Information for**
### CHROMOMETHYLTRANSFERASE3/ KRYPTONITE maintain the *sulfurea* paramutation in *Solanum lycopersicum*

Cláudia Martinho, Zhengming Wang, Andrea Ghigi, Sarah Buddle, Felix Barbour, Antonia Yarur, Quentin Gouil, Sebastian Müller, Maike Stam, Chang Liu and David C. Baulcombe

David C. Baulcombe


**This PDF file includes:**

Supplementary text
Figures S1 to S13
Tables S1 to S5
SI References

#### Material and Methods

**Plant methods.** Plants were germinated and grown in *F2* compost and transferred to *John Innes 2* compost three weeks after germination. Plants were grown in 16h light (22°C) and 8h dark (18°C) cycles with 70% humidity and light intensity 300 µmol lx $m^{-2}$ x $s^{-1}$ PAR**.** Emasculation and pollination were carried out following the Tomato Genetics Resource Centre guidelines - https://tgrc.ucdavis.edu/Guidelines_Emasculating_and_Pollinating_Tomatoes.pdf. The plant lines used for this study are summarised in **Table S2**. For molecular analyses leaves were excised from 4-week-old plants using sharp tweezers. In the case of *sulf* plants, leaves with higher degree of chlorosis were selected for each individual. Tissue was flash frozen in liquid nitrogen.

**Genotyping.** Genomic DNA was isolated from 100mg of leaf tissue from 4-week-old plants using the Dneasy Plant Mini Kit (Qiagen) according to the manufacturer's instructions. For all genotyping reactions genomic DNA was diluted to a final concentration of 10ng/µL in nuclease free water. *cmt3* and *kyp* mutants were genotyped using oligonucleotides as described in (1) and also listed in **Table S3**. The mutant *cmt3* allele was amplified using Phusion High-Fidelity DNA Polymerase (Thermo Scientific). The WT CMT3 alleles and KYP WT and mutant alleles were amplified with DreamTaq DNA Polymerase (Thermo Scientific). *nrpe1* mutants were genotyped with the oligonucleotides listed in **Table S3** followed by T7 endonuclease digestion. Two digestion reactions were setup per sample: one with 10µL of sample PCR reaction alone and the other with 5µL of sample PCR reaction mixed together with 5µL of a WT cv. M82 PCR reaction to allow identification of *nrpe1* homozygotes upon T7 digestion. 1.5µL Buffer 2 (New England Biolabs) and 1.5µL of nuclease free water were added to each reaction and denatured for 5 min at 95°C, followed by annealing with a 95°C – 85°C ramp with 2°C/sec increments and a ramp 85°C – 25°C with 0.1°C/sec increments. 2µL of T7 endonuclease (New England Biolabs) diluted to 2U/µL in 1X Buffer 2 (New England Biolabs) were added to each reaction and incubated at 37°C for 1 hour. The fragments were separated in a 1.5% agarose gel immediately after digestion. All genotyping PCR conditions are detailed in **Table S4**.

**Tomato CHG subcontext analysis**. Whole-genome bisulphite sequencing data for duplicates of $TAB2^{+}$ and $TAB2^{sulf}$ epigenotypes were retrieved from BioProject SRP066362 (2). Paired reads were trimmed with Trim Galore! v0.4.4 (https://www.bioinformatics.babraham.ac.uk/projects/trim_galore/) and aligned to *Solanum lycopersicum* assembly Heinz 1706 version 3.00 with Bismark v0.19.1 (3). Following deduplication and methylation report generation with Bismark, differentially methylated regions were defined using HOME v1.1 (4), filtering for at least 20% difference in absolute methylation levels between the TAB2+ and TAB2sulf plants, and at least 5 cytosines in the DMR. Extracted data was used for subcontext analysis using an in-house R script (**Supplemental file S1**). The output was parsed and plotted using the scripts available at https://github.com/claudiamartinho/Martinhoetal2021.

**McrBC-qPCR.** Genomic DNA was isolated from 100mg of leaf tissue using Dneasy Plant Mini Kit (Qiagen) according to the manufacturer's instructions. The DNA samples used for McrBC digestion were the same as the samples using for genotyping. To determine the proportion of DNA methylation at DMR1 McrBC digestion were carried out followed by quantitative real time PCR (qPCR). McrBC digestion was performed as described in (5). 10µl qPCR reactions were assembled using 4µg of digested or undigested genomic DNA and 1X Luna® Universal qPCR Master Mix (New England Biolabs). qPCR was carried out with one denaturation step at 95°C for 3 min, followed by 40 cycles of denaturation at 95°C for 10 sec annealing at 60°C for 20 sec, and primer extension at 72°C for 30 sec. Upon completion of the cycling steps, a final extension at 72°C for 5 min on a CFX384 system (Bio-Rad) was performed. The sequences of oligonucleotides used for McrBC-qPCR are listed in **Table S3**. DNA methylation proportion was calculated using the formula $100 \times \{1-2^{(\mathrm{Ct\ undigested} - \mathrm{Ct\ digested})}\}$. Negative methylation values were converted to 0% DNA methylation as they represent no loss of amplification and therefore no methylation.

**Expression analysis.** Total RNA isolation was performed with the Direct-zol RNA Miniprep (Zymo Research) and Trizol reagent (Invitrogen) according to the manufacturer's instructions. For quantitative RT-PCR analyses 1µg of total RNA was reverse transcribed using RevertAid First Strand cDNA Synthesis Kit (Thermo Scientific) according to the manufacturer's instructions with random hexamer primers. Quantitative PCR was performed in reactions containing 1X Luna® Universal qPCR Master Mix (New England Biolabs) on a CFX384 system (Bio-Rad). qPCR was carried out denaturation step at 95°C for 3 min, followed by 40 cycles of denaturation at 95°C for 10 sec, annealing/extension at 60°C for 30 sec. Relative expression was calculated using the ΔΔct method (2−ΔΔct) using the geometric mean of two reference genes **(Table S3)**. Oligonucleotides used for expression analyses are listed in **Table S3**.

**sRNA-seq.** Total RNA isolation was performed with the Direct-zol RNA Miniprep (Zymo Research) and Trizol reagent (Invitrogen) according to the manufacturer's instructions in leaf tissue of 4-week-old plants. For M82/*sulf* and F2 CMT3 sets, sRNA libraries were prepared using the NEBNext multiplex small RNA library prep kit (New England Biolabs) according to the manufacturer's instructions. Libraries were indexed during the PCR step with 12 cycles and size-selected using BluePippin. Pooled libraries were sequenced on a NextSeq500 (Illumina). NRPE1 library preparation and sequencing were outsourced to Novogene. Raw data (already demultiplexed and available in fastq format) was processed using the Snakemake pipeline available at https://github.com/seb-mueller/snakemake_sRNAseq. Briefly, raw data was quality controlled using FastQC (v0.11.7) followed by 3' adaptor removal (trimming) using cutadapt removing Illumina universal adapters. All sequences <15 nt and >40 nt in length were discarded, and the remaining sequences mapped to the reference genome (Heinz 1706 genome version 3.0). Mapping was performed using Bowtie version 1.2 with uniquely mapping with "bowtie --wrapper basic-0 -v 0 -k 1 -m 1 --best –q" which only reports sRNAs mapping to unique locations. 0 mismatches were employed using bowtie version 1.2. DMR1 sRNA quantity was normalized as count per million (CPM) basing it on the total number of reads mapped at DMR1 (please see coordinates in Table S1) using deeptools version 3.3.1. The config.yaml file used for this analysis is supplied as **Supplemental file S2**.

**Whole genome bisulphite sequencing.** Genomic DNA was isolated from 100 mg leaf tissue 4-week-old plants using Dneasy Plant Mini Kit (Qiagen). Library preparation and sequencing were carried out by Novogene. In brief, DNA samples were fragmented into 200-400bp using Covaris S220. Terminal repairing, A-ligation, methylation sequencing adapters ligation were performed to the DNA fragments. Bisulphite treatment was carried out with Accel-NGS Methyl-Seq DNA Library Kit (IIllumina Cat No. 30096) followed by size selection and PCR amplification steps. Whole genome bisulphite processing raw data (paired-end; already demultiplexed and available in fastq format) was processed using the bisulfite Snakemake pipeline available at https://github.com/seb-mueller/snakemake-bisulfite. Briefly, raw data was quality controlled using FastQC (v0.11.7) followed by 3' adaptor removal (trimming) using trim_galore (https://www.bioinformatics.babraham.ac.uk/projects/trim_galore/) removing Illumina universal adapters. Mapping and cytosine methylation calling was done using Bismark (3) on the reference genome (Heinz 1706 genome version 3.00) in CpG, CHG and CHH contexts. The config.yaml file used for this analysis is supplied as **Supplemental file S3**.

**ChIP-qPCR.** Chromatin extraction was carried out as described in (1) using 1g of leaf tissue of 4 week-old plants. Chromatin was fragmented between 200-600bp in a Covaris E220 evolution (duty cycle: 20%, peak intensity: 140, cycles of burst: 200, time: 3 min). Immunoprecipitation was carried out as described in (1) with Anti-Histone H3 (di methyl K9) antibody - ChIP Grade (abcam, ab1220) and Anti-Histone H3 (tri methyl K4) antibody - ChIP Grade (abcam, ab8580). The material was reverse cross-linked by adding NaCl to a final concentration of 200mM and incubating at 65°C O/N. This was followed by a 30 min treatment with 1µL RNase A (Thermo Scientific) at 37°C and a 90 min treatment with 1.5 µl Proteinase K (Thermo Scientific) at 65°C. DNA was purified using the MinElute Kit (Qiagen) according to manufacturer's instructions and eluted in 35µl EB buffer. The eluted DNA was used to quantify enriched DNA fragments by standard qPCR methods using 1X Luna® Universal qPCR Master Mix (New England Biolabs) on a CFX384 system (Bio-Rad). Enrichment of DNA fragments for H3K9me2 and H3K4me3 analysis were calculated as % input (2(Ct input adjusted – Ct IP)*100) and normalised using the enrichment found for an unrelated reference locus *CAC3*. Oligonucleotides employed in this analysis are listed in **Table S3.**

**Hi-C.** 0.5g leaf tissue of 4-week-old plants (per replicate) was fixed in 1% formaldehyde (v/v) and 0.01% Triton-X (v/v) for 20 min using vacuum infiltration. The reaction was quenched by adding glycine to a final concentration of 125 mM for 10 min using vacuum infiltration. The leaf tissue was rinsed 3 times with MilliQ water, pat dried and flash frozen in liquid Nitrogen. The *in situ* Hi-C library preparation was performed essentially as described in (6). For individual samples in each replicate was homogenized for preparing libraries. The libraries were sequenced on an Illumina NextSeq500 instrument with 2 x 75 bp reads. The Hi-C raw reads were mapped to *Solanum lycopersicum* genome Heinz 1706 assembly SL3.00 with an iterative mapping pipeline (6). The removal of PCR duplicates and reads filtering were performed as described in (6). Hi-C reads of each sample are summarised in **Table S5**. Hi-C map normalization with bin size 100kb was done using the "HiTC" package in R (7).

**Maize CHG subcontext analysis**. For bisulphite sequencing, genomic maize DNA was extracted (8) from pools of 30-100 embryos or half of an ear. Bisulphite treatment was performed using 400 ng of DNA and the EZ DNA Methylation-Gold kit (Zymo Research, D5006). The DNA regions of interest were PCR amplified (10 min 95°C, followed by 40 PCR cycles (30 sec 95°C, 30 sec appropriate annealing temp, 30 sec 72°C), and 5 min at 72°C). The PCR was performed using MethylTaq DNA polymerase (Diagenode, C09010010). For primer sequences, amplicon sizes and annealing temperatures see **Table S5**. In order to monitor for complete bisulphite conversion, a conversion control (Fie2 fragment) was amplified and analysed in each experiment **(Fig S13)** (similar to the -302 to -91 fragment described in (9)). PCR fragments were ligated into the pJET 1.2 vector (CloneJet PCR Cloning Kit, Thermo Scientific) according manufacturer's instructions. Positive Clones were identified by colony-PCR, Plasmid DNA was isolated from positive clones (GeneJet Plasmid Miniprep Kit, Thermo scientific) and subjected to Sanger sequencing. For the conversion control 14-16 clones were analysed and for each other fragment 22-31 clones were analysed. Frequency of DNA methylation at individual CHG motifs was inferred using Kismeth (10).

**Data visualisation and statistical analysis**

Plots for ChIP-seq, sRNA-seq and subcontext analysis were carried out using ggplot2 (11) and computationally reproducible scripts are available at https://github.com/claudiamartinho/Martinhoetal2021. Plots depicting DNA methylation levels determined by McRBC and expression levels were generated by employing the webtool PlotsOfData (12). Statistical tests were carried out in R using the function *wilcox.test* and *p.adjust.method = "BH"*. Genome browser images were generated in IGV v2.7.2 (13).

**Data availability**

All sequencing datasets, including sRNA, bisulphite and Hi-C were deposited at ArrayExpress (EMBL-EBI) under the accession numbers: E-MTAB-10556, E-MTAB-10557, E-MTAB-10565, E-MTAB-10568 and E-MTAB-10574.

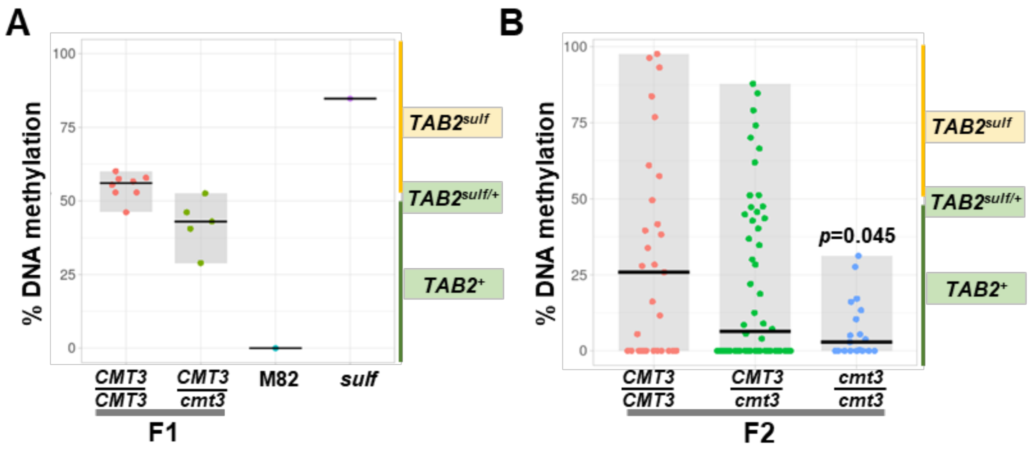


**Fig. S1.** – **DMR1 DNA methylation is reduced in *cmt3* mutants.** Jittered dots depict DNA Methylation percentage at DMR1 in individual plants determined by McrBC-qPCR. The summary of the data is shown as horizontal line indicating the median. Grey boxes illustrate the data range. Plants result from the cross between *CMT3/CMT3 TAB2*$^{sulf}$ and *CMT3/cmt3 TAB2*$^{+}$ (Fig 1C). **A–** F1 sibling plants: *CMT3/CMT3* n=8; *CMT3/cmt3* n=5; controls: M82- (*S. lycopersicum cv. M82) CMT3/ CMT3 TAB2*$^{+}$ *n=1; sulf* - (*S. lycopersicum cv. Lukullus) CMT3/CMT3 TAB2*$^{sulf}$ *n=1*. **B–** F2 sibling plants: *CMT3/CMT3* n= 29; *CMT3/cmt3* n=52; *cmt3/cmt3* n=19. p-value *cmt3/cmt3* versus *CMT3/CMT3* was calculated employing a Mann-Whitney-Wilcoxon test. Yellow boxes refer to plants displaying *sulf* chlorosis and green boxes refer to green plants.

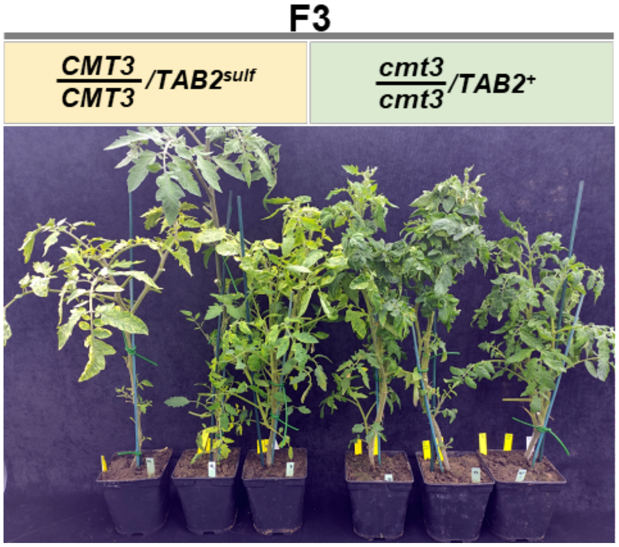


**Fig. S2. CMT3 suppresses *sulf* chlorosis**. 2-month-old F3 sibling plants *CMT3/CMT3 TAB2*$^{sulf}$ and *cmt3/cmt3 TAB2*$^{+}$. Yellow boxes refer to plants displaying *sulf* chlorosis and green boxes refer to green plants.

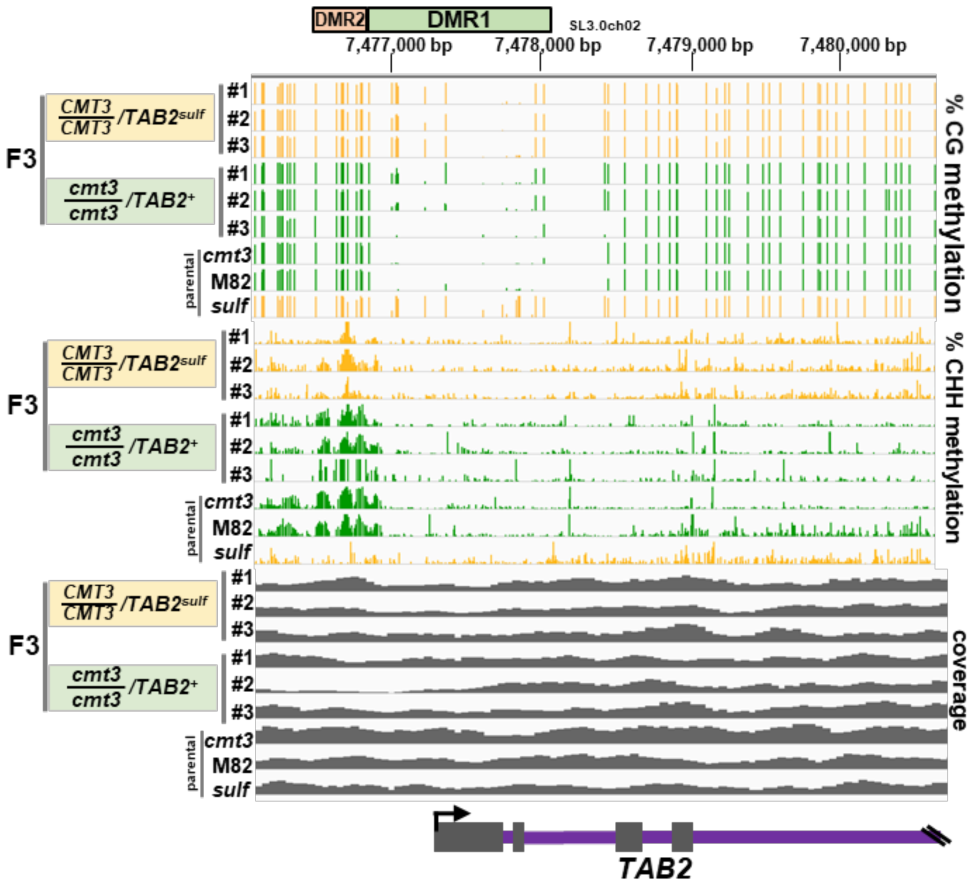


**Fig. S3.** – CG and CHH DNA methylation levels at DMR1 in F3 plants (F3 Fig. 1). *TAB2* IVG screenshot bisulphite sequencing data, % CG DNA methylation, range [0-100]. % CHH DNA methylation, range [0-100]. Green tracks refer to green leaf phenotype and yellow tracks refer to plants which display chlorosis. Sequencing coverage range [0-65]. Yellow tracks and boxes refer to plants displaying *sulf* chlorosis and green tracks and boxes refer to green plants.

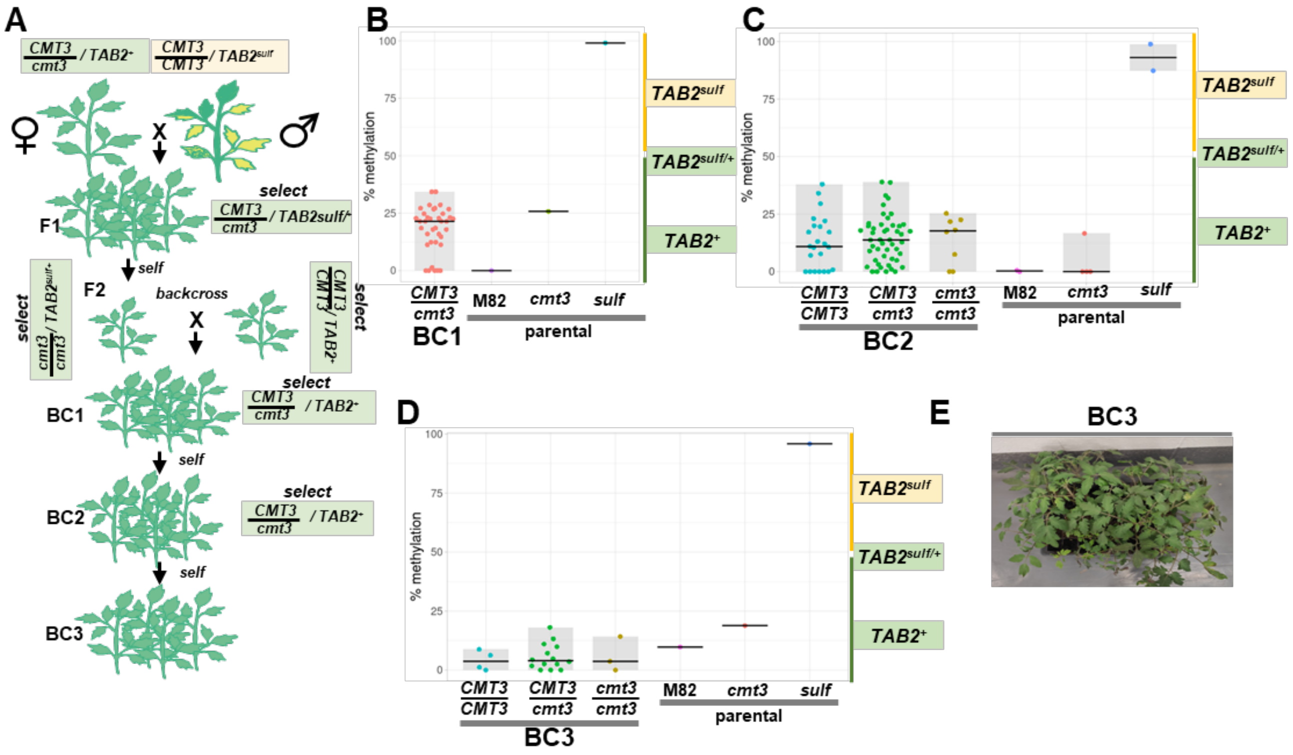


**Fig. S4. CMT3 maintains *sulf* memory. A–** Diagram illustrates crossing scheme used to obtain backcrossed populations. **B–D** Jittered dots depict % of DNA methylation at DMR1 in individual plants determined by McrBC digestion followed by qPCR. The summary of the data is shown as horizontal line indicating the median. Grey boxes illustrate the data range. **B–** Plants denote first backcross generation (BC1 - Fig. S4A) n=21; Control plants: M82- (*S. lycopersicum cv. M82) CMT3/CMT3* $TAB2^{+}$ *n=1; sulf-* (*S. lycopersicum cv. Lukullus) CMT3/CMT3* $TAB2^{sulf}$ *n=1; F2 parent - cmt3/cmt3* $TAB2^{+}$ n=1. **C–** Second backcross generation sampling (BC2 – Fig. S4A). BC2 plants: *CMT3/CMT3* $TAB2^{+}$ n=23; *CMT3/cmt3* $TAB2^{+}$ n=48; *cmt3/cmt3* $TAB2^{+}$ n=8. Control plants (these samples were processed in different batches and one control plant was added per batch). M82- (*S. lycopersicum cv. M82) CMT3/CMT3* $TAB2^{+}$ *n=2; sulf-* (*S. lycopersicum cv. Lukullus) CMT3/CMT3* $TAB2^{sulf}$ n=2*; cmt3-cmt3/cmt3* $TAB2^{+}$ n=4. **D–**Third backcross generation sampling (BC3 – Fig. S4A). BC3 plants: *CMT3/CMT3* $TAB2^{+}$ n=3; *CMT3/cmt3* $TAB2^{+}$ n=14; *cmt3/cmt3* $TAB2^{+}$ n=3. Control plants: M82- (*S. lycopersicum cv. M82) CMT3/CMT3* $TAB2^{+}$ n=1*; sulf-* (*S. lycopersicum cv. Lukullus) CMT3/CMT3* $TAB2^{sulf}$ n=1*; cmt3- cmt3/cmt3* $TAB2^{+}$ n=1. **F–** BC3 population consists of 100% green plants – 4-week-old plants. Yellow boxes refer to plants displaying *sulf* chlorosis and green boxes refer to green plants.

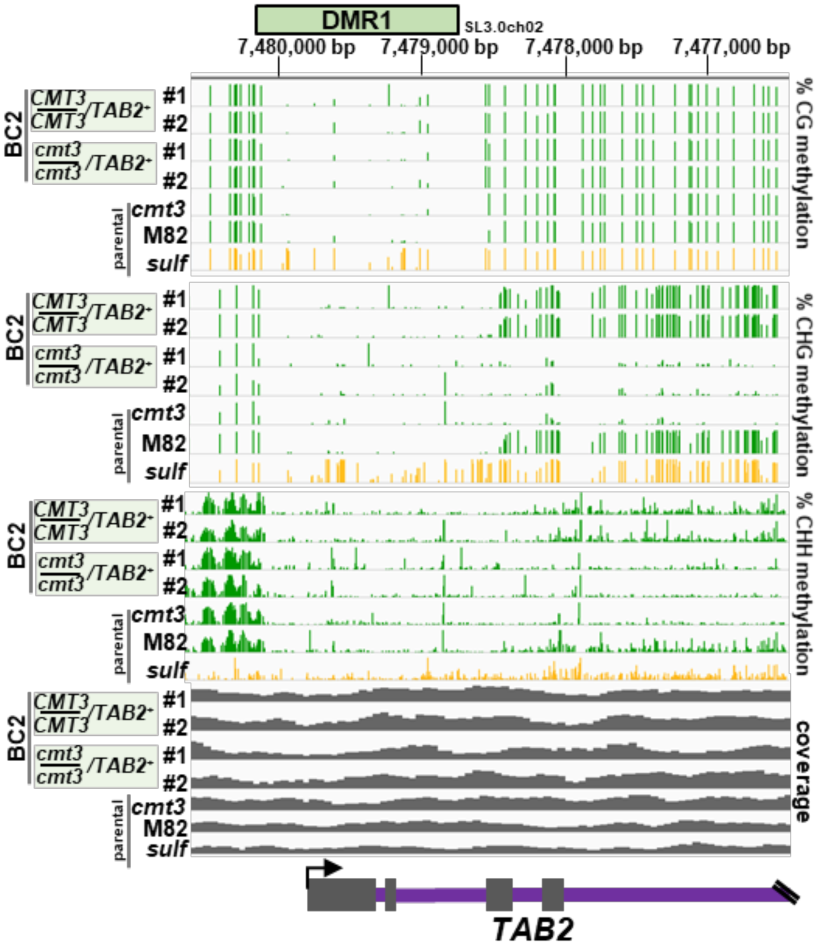


**Fig. S5. Backcrossed populations don't recover DMR1 DNA methylation.** CG and CHH DNA methylation levels at DMR1 in backcrossed plants – generation 2 (BC2) plants (Fig. S4A). *TAB2* IGV screenshot bisulphite sequencing data, % CG DNA methylation, range [0-100]. % CHH DNA methylation, range [0-100]. Green tracks refer to green leaf phenotype and yellow tracks refer to plants which display chlorosis. Sequencing coverage range [0-72]. Yellow tracks refer to plants displaying *sulf* chlorosis and green boxes and tracks refer to green plants.

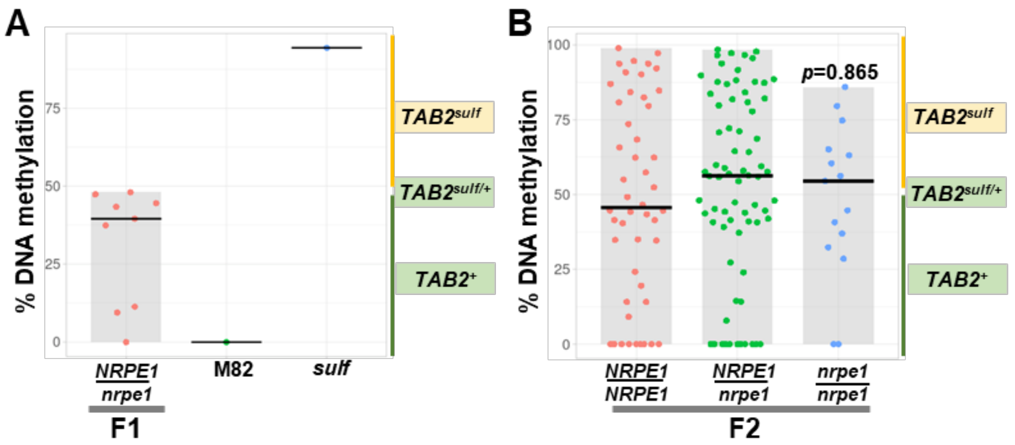


**Fig. S6. DMR1 DNA methylation is unaffected in *nrpe1* mutants.** Jittered dots depict DNA methylation percentage at DMR1 in individual plants determined by McrBC-qPCR. The summary of the data is shown as horizontal line indicating the median. Grey boxes illustrate the overall data range. Plants are derived from crosses between *NRPE1/NRPE1 TAB2$^{sulf}$* and *nrpe1/nrpe1 TAB2$^{+}$* (Fig. 2A). **A–** F1 sibling plants: *NRPE1/nrpe1* n=9; controls: M82- (*S. lycopersicum cv. M82) CMT3/ CMT3 TAB2$^{+}$ n=1; sulf* - (*S. lycopersicum cv. Lukullus) CMT3/CMT3 TAB2$^{sulf}$ n=1*. **B–** F2 sibling plants: *NRPE1/NRPE1* n= 48; *NRPE1/ nrpe1* n=72; *nrpe1/nrpe1* n=15. *p-value nrpe1/nrpe1* versus *NRPE1/NRPE1* was calculated by employing a Mann-Whitney-Wilcoxon test. Yellow boxes refer to plants displaying *sulf* chlorosis and green boxes refer to green plants.

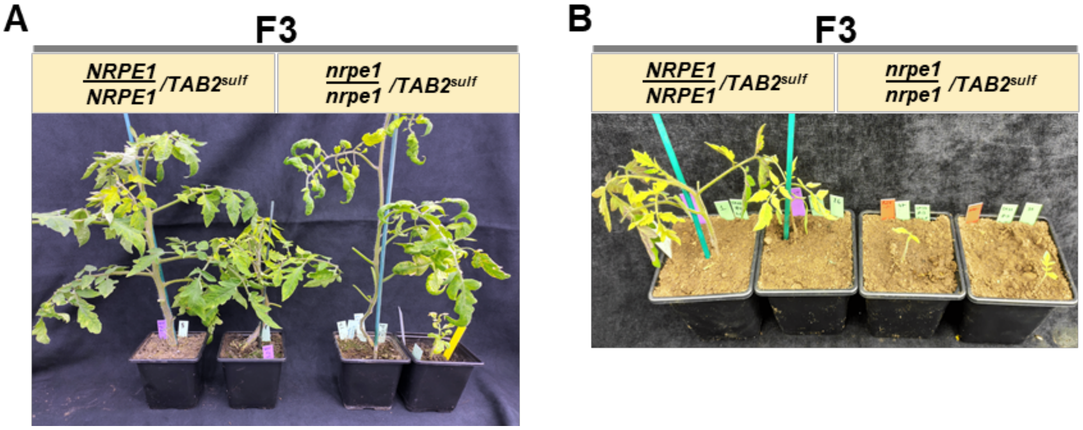


**Fig. S7. *Sulf* phenotype is maintained in the *nrpe1* background.** F3 plants derived from crossing *NRPE1/NRPE1* $TAB2^{sulf}$ and *nrpe1/nrpe1* $TAB2^{+}$ (Fig. 2A). **A–** 6-week-old. **B–** 4-week-old. Yellow boxes refer to plants displaying *sulf* chlorosis.

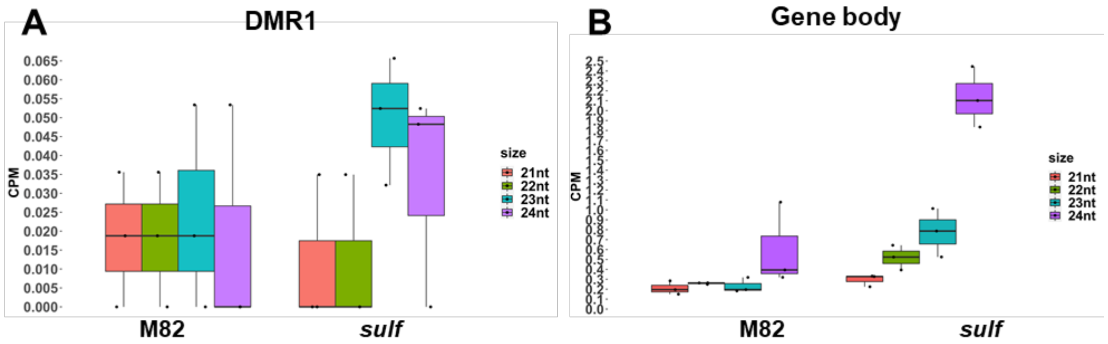


**Fig. S8. sRNA accumulation in *sulf* at TAB2.** sRNA accumulation in count per million (CPM). Jitter dots represent biological replicates n=3. The summary of the data is shown as horizontal line indicating the median. Error bars represent standard deviation. **M82–** (*S. lycopersicum cv. M82*) *TAB2*$^{+}$ n=3; ***sulf*–** (*S. lycopersicum cv. Lukullus*) *TAB2*$^{sulf}$ n=3. **A–** DMR1. **B–** Gene body. Coordinates are provided in Table S1.

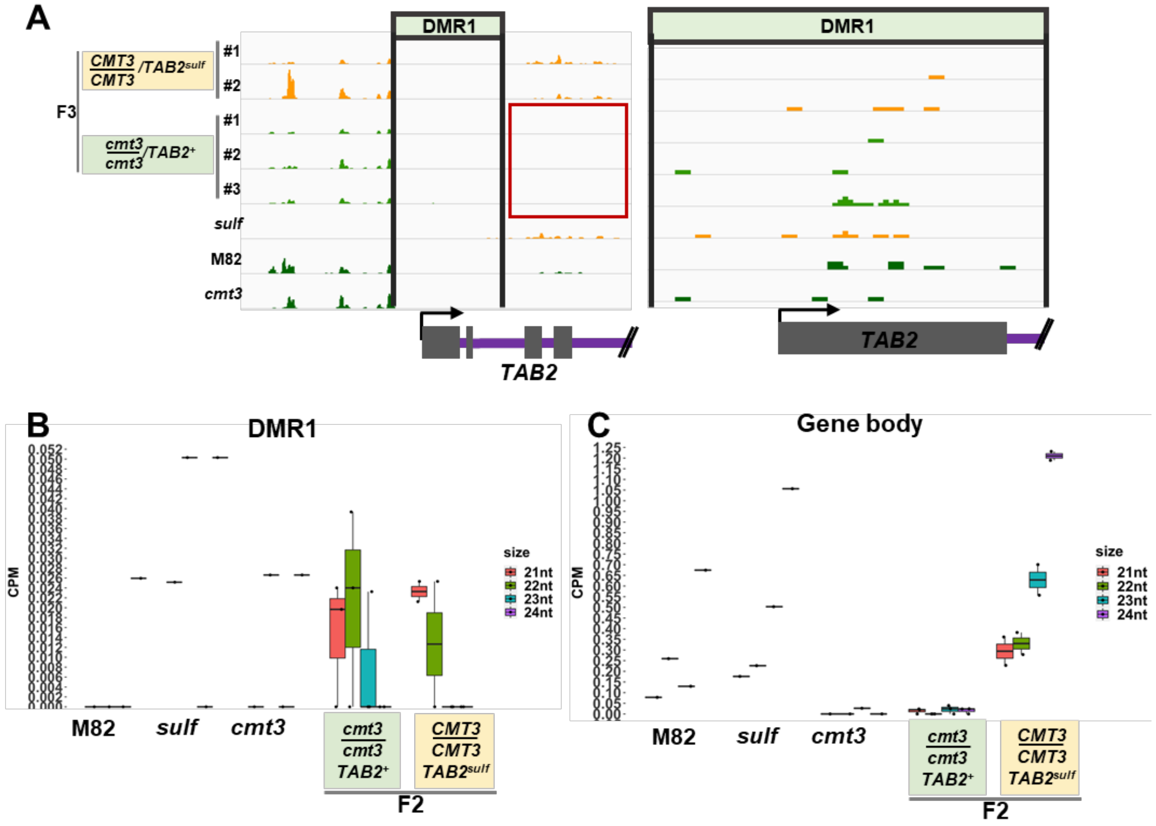


**Fig. S9. sRNA accumulation at *TAB2* in *cmt3* and *CMT3* F2 plants** (Fig. 1C and Fig. S1 B). **A–** TAB2 IGV screenshot small-RNA sequencing data. Left panel – sRNA accumulation in DMR1 and TAB2 gene body, range [0-8.53]; Right panel – zoom in DMR1 region, range [0-8.53]. Red box highlights absence of sRNA accumulation in the gene body region in the *cmt3* background. **B** and **C**– Boxplots depict siRNA accumulation in counts per million (CPM). Jitter dots represent biological replicates n=3. The summary of the data is shown as horizontal line indicating the median. Error bars represent standard deviation. **B–** DMR1. **C–** Gene body. **A-C**: controls: **M82–** (*S. lycopersicum cv. M82*) $TAB2^{+}$ n=3; ***sulf*–** (*S. lycopersicum cv. Lukullus*) $TAB2^{sulf}$ n=3; ***cmt3*–** control *cmt3/cmt3* $TAB2^{+}$ (*S. lycopersicum cv. M82*) F3 *CMT3/CMT3* $TAB2^{+}$ n= 2; F2 plants: *cmt3/cmt3* $TAB2^{+}$ n=3, *CMT3/CMT3* $TAB2^{sulf}$ n=2. Yellow boxes refer to plants displaying *sulf* chlorosis and green boxes refer to green plants.

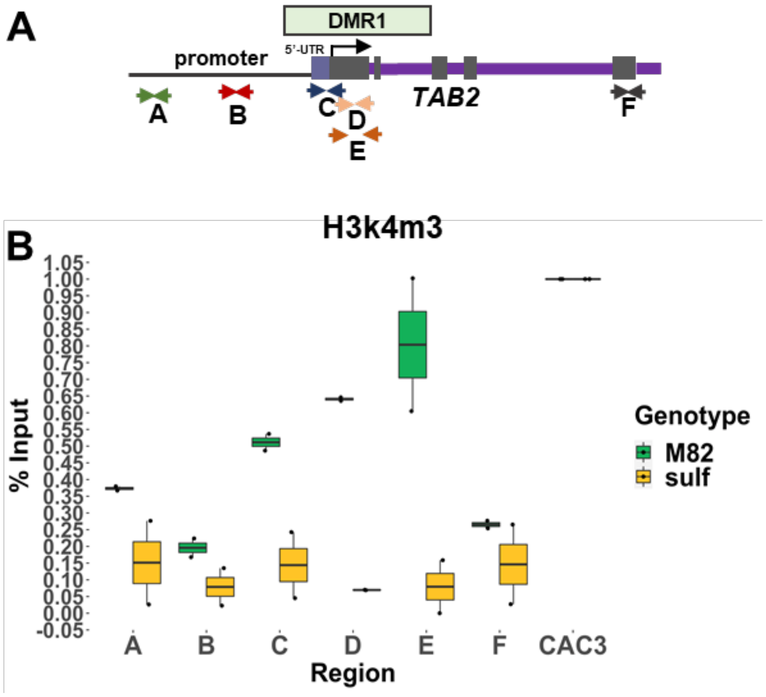


**Fig. S10. H3K4me3 levels at *TAB2*. A–** Diagram represents the relative oligonucleotide position spanning the *TAB2* locus used for ChIP-qPCR experiment in Fig. S10B (sequences are listed in Table S3). **B–** Box plot depicts H3k4m3 enrichment per % input normalised to *CAC3* reference locus determined by Chip-qPCR. Jittered dots represent different biological replicates. **M82–** *S. lycopersicum* cv. M82 *TAB2*$^{+}$ (green) n= 2; ***sulf*–** *S. lycopersicum cv. Lukullus TAB2*$^{sulf}$ (yellow) n=2. The summary of the data is shown as horizontal line indicating the median of biological replicates. Error bars represent the standard deviation.

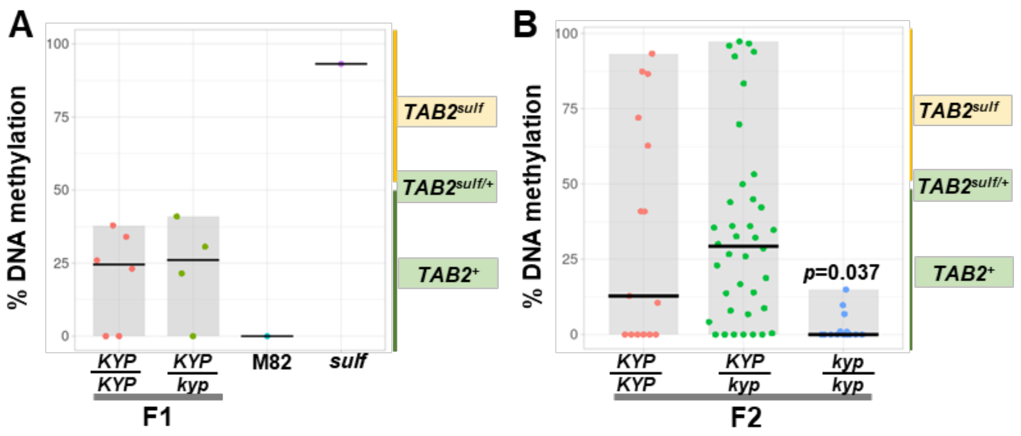


**Fig. S11. DMR1 DNA methylation is reduced in *kyp* mutants.** Jittered dots depict DMR1 DNA methylation percentage in individual plants determined by McrBC-qPCR. The summary of the data is shown as horizontal line indicating the median. Grey boxes illustrate the data range. Plants result from the cross between *KYP/KYP* $TAB2^{sulf}$ and *KYP/kyp* $TAB2^{+}$ (Fig. 3C). **A–** F1 sibling plants: *KYP/ KYP* n=8; *KYP /kyp* n=5; controls: **M82–** (*S. lycopersicum cv. M82*) $TAB2^{+}$ n=3; ***sulf*–** (*S. lycopersicum cv. Lukullus*) $TAB2^{sulf}$ n=3. **B–** F2 sibling plants: *KYP/KYP* n=15; *KYP/kyp* $TAB2^{sulf}$ n=38; *kyp/kyp* $TAB2^{+}$ n=15. p-value *kyp/kyp* versus *KYP/KYP* was calculated employing a Mann-Whitney-Wilcoxon test. Yellow boxes refer to plants displaying *sulf* chlorosis and green boxes refer to green plants.

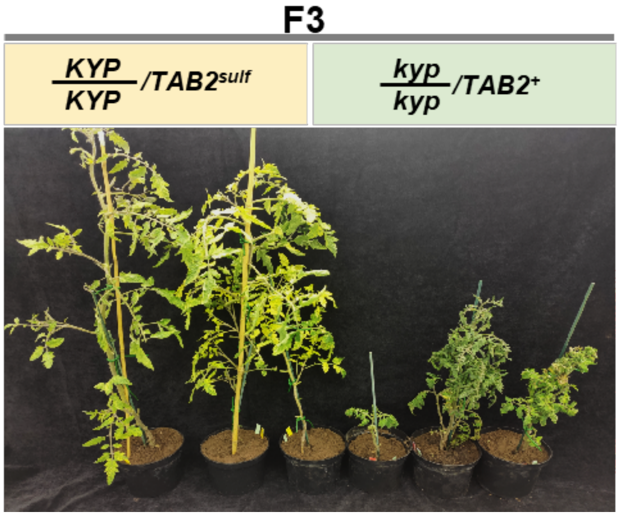


**Fig. S12. KYP suppresses *sulf* chlorosis**. 2-month-old F3 sibling plants *KYP/KYP TAB2$^{sulf}$* and *kyp/kypTAB2$^{+}$*. Yellow boxes refer to plants displaying *sulf* chlorosis and green boxes refer to green plants.

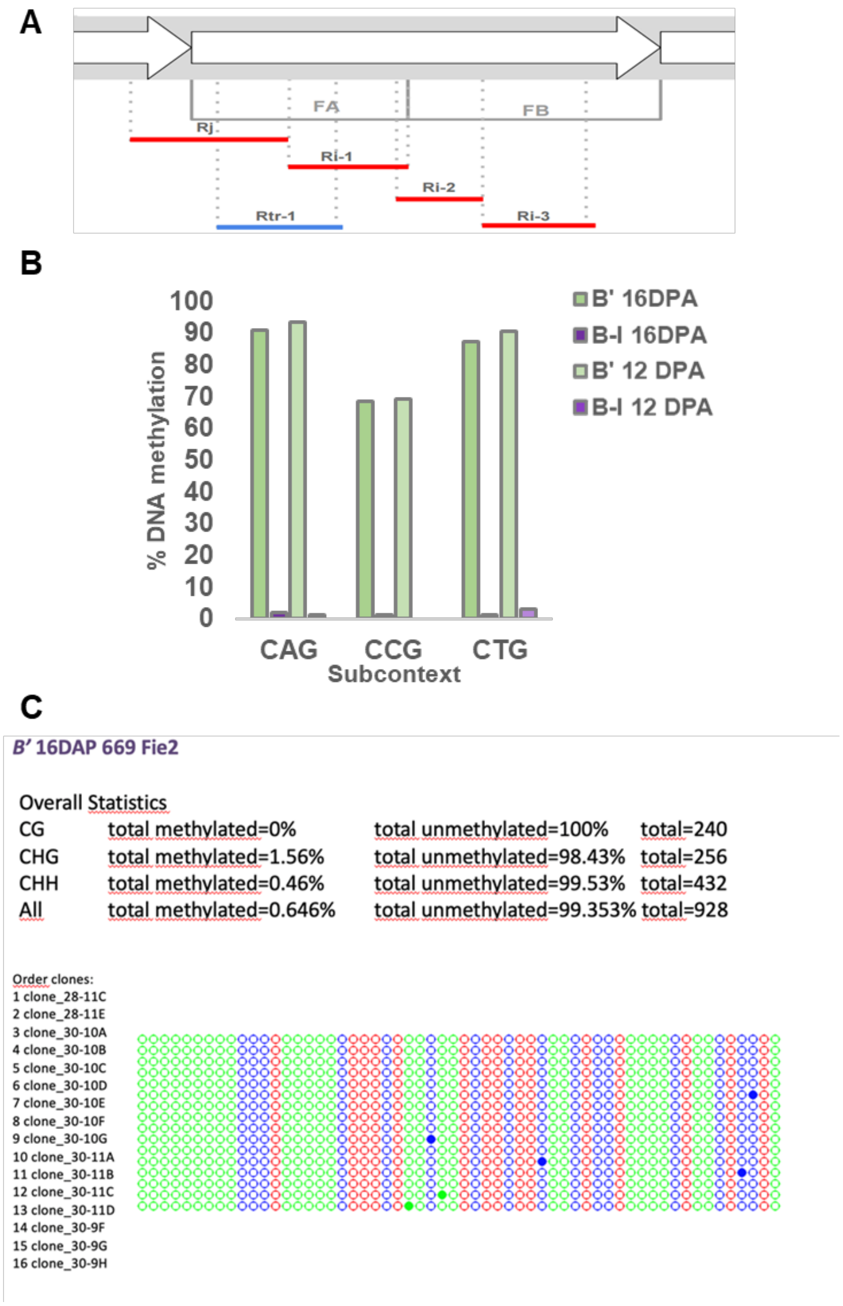


**Fig. S13. CHG subcontext analysis at the *b1* locus in maize**. **A –** Schematic depiction of one repeat of the b1 hepta-repeat. One repeat is 853 bp long. For Bisulfite-sequencing fragment Rj (320 bp) was analysed. **B–** Mean percentage of DNA methylation at the different CHG subcontexts at Rj region. n CAG = 7; n CCG = 4; n CTG = 7; **C–** Average and clone-distribution of DNA methylation levels at the unmethylated *Fie2* locus in B' samples – low methylation levels confirm bisulphite conversion was complete.

| Name | Chromosome | start | end |
|---|---|---|---|
| **Control 1** | SL3.00ch02 | 3606037 | 3607183 |
| **Control 2** | SL3.00ch02 | 32797030 | 32798176 |
| **Control 3** | SL3.00ch02 | 35314142 | 35315288 |
| **Control 4** | SL3.00ch02 | 35727400 | 35728546 |
| **Control 5** | SL3.00ch02 | 44030115 | 44031261 |
| **DMR1** | SL3.00ch02 | 7478826 | 7480096 |
| **Gene body** | SL3.00ch02 | 7477602 | 7478748 |

**Table S1.** DMR1 and Gene body coordinates used for the analyses in this manuscript. Random control regions in SL3.00ch02 were used for subcontext analysis Fig. 1B.

| Plant line | Epigenotype | Genotype | | | | Described in |
|---|---|---|---|---|---|---|
| | ***TAB2* (Solyc05g005200)** | ***CMT3* (Solyc01g006100)** | ***KYP* (Solyc02g094520)** | ***NRPE1* (Solyc01g096390)** | **cultivar** | |
| ***sulfurea (sulf)*** | *TAB2sulf* | WT | WT | WT | Lukullus | (14, 15) |
| **M82** | *TAB2+* | WT | WT | WT | M82 | **TGRC - acession LA3475** |
| ***cmt3*** | *TAB2+* | CRISPR deletion | WT | WT | M82 | (1) |
| ***kyp*** | *TAB2+* | WT | CRISPR deletion | WT | M82 | (1) |
| ***nrpe1*** | *TAB2+* | WT | WT | CRISPR deletion | M82 | (16) |

**Table S2.** Plant lines employed in this study.

| Number | Name | Gene ID | Sequence 5'-3' | orientation | Description | Purpose |
|---|---|---|---|---|---|---|
| **204** | 204_CMT3_WT | Solyc01g006100 | TAGGGTGATGTTGATGTTGTATG | Forward | Genotyping *cmt3* mutant | **Genotyping** |
| **205** | 205_CMT3_MU | | TAGGGTGATGTTGATGCGGT | Forward | | |
| **206** | 206_CMT3_R | | ATCGCCCAAGGAAACCATTG | Reverse | | |
| **201** | 201_KYP_WT | Solyc02g094520 | AAGATTTAGCAGAAACCTGCA | Forward | Genotyping *kyp* mutant | |
| **202** | 202_KYP_MU | | AAGATTTAGCAGAAACAGAGG | Forward | | |
| **203** | 203_KYP_R | | TTGGCTGTTGCAGTTTTCCCTGAG | Reverse | | |
| **275** | nrpe1 Fw | Solyc01g096390 | AGATGAAGACCTTGTTTATCAG | Forward | Genotyping *nrpe1* mutant | |
| **276** | nrpe RV | | TGTGATAACATCATTCAGAATTG | Reverse | | |
| **23** | 23_Small_fw | Solyc05g005200 | TTTGATGGCTTGTCTAAGCTGC | Forward | DMR1 McrBC | **McrBC-qPCR** |
| **24** | 24_Small_rv | | AGCTCTCAGGATCAATTTCAGGG | Reverse | | |
| **35** | CAC3 fw qPCR | Solyc08g006960 | TGTCGTCAAAAGGTAGCGTCCTCCGT | Forward | Reference genes | **Expression analysis** |
| **36** | CAC3 rv qPCR | | AGCCCAACTTCAGATCAGGCATTCCA | Reverse | | |
| **37** | SKP1 FW qPCR | Solyc11g042930 | TCAGGGCACCCTCTTCGATCTGATCCT | Forward | | |
| **38** | SKP1 RV qPCR | | TGTCTGCCACGGTCTGGCAAGTGA | Reverse | | |
| **43** | SLTAB2_qPCR_CM1 | Solyc05g005200 | AAGGAGCCAAGCCACTTCTT | Forward | TAB2 expression | |
| **44** | SLTAB2_qPCR_CM1 | | GCAACTGGACAAAAGCCCAC | Reverse | | |
| **73** | TAB2_Chip-qPCR3_fw | upstream Solyc05g005200 | GCCTCACACTCTTCTTTGCC | Forward | Region A | **ChIP-qPCR** |
| **74** | TAB2_Chip-qPCR4_rev | | GTTGCCAGCTAGCTCACAAA | Reverse | | |
| **232** | 232TAB2_mnase_fw11 | upstream Solyc05g005200 | AAAGTGAGAGATGCGAAAGAGAGA | Forward | Region B | |
| **233** | 233TAB2_mnase_rev12 | | TTCGTAGCAAACAATTGTCATTCA | Reverse | | |
| **238** | 238TAB2_mnase_fw17 | upstream Solyc05g005200 | CGATATTTAGGCCGAAGAAAGAG | Forward | Region C | |
| **239** | 239TAB2_mnase_rev18 | | ACAAGCCATCAAACCCAAATT | Reverse | | |
| **236** | 236TAB2_mnase_fw15 | upstream Solyc05g005200 | AATTTGGGTTTGATGGCTTGT | Forward | Region D | |
| **237** | 237TAB2_mnase_rev16 | | TTGCAACTGATTTAAAGGGGAAT | Reverse | | |
| **23** | 23_Small_fw | | TTTGATGGCTTGTCTAAGCTGC | Forward | Region E | |
| **24** | 24_Small_rv | | AGCTCTCAGGATCAATTTCAGGG | Reverse | | |
| **77** | TAB2_Chip-qPCR7_fw | | GGAAGCAGCAAAGAAAGCCT | Forward | Region F | |
| **78** | TAB2_Chip-qPCR8_rev | | AGGAGGTGGCAGGTTTAACA | Reverse | | |
| **79** | CAC3_Chip-qPCR1_fw | Solyc08g006960 | GATGAGTGACGGAGCCAGTA | Forward | CAC3 used for normalisation | |
| **80** | CAC3_Chip-qPCR2_rev | | CACAAAAGAGGCCTGCAGAG | Reverse | | |
| **207** | 207_T135_K9control_F | - | CCAGCCATAACAACCAACTTC | Forward | Amplifies T135 transposon elements | |
| **208** | 208_T135_K9control_R | - | GCAGACCACCAAATCCAACTC | Reverse | | |

**Table S3.** Oligonucleotides used for tomato experiments – all oligonucleotides were designed using the *Solanum lycopersicum* assembly Heinz 1706 version SL3.00.

**WT *CMT3* allele**

| Component | volume (µL) |
|---|---|
| Green buffer (10X) | 2 |
| dNTPs (10mM each) | 0.5 |
| Oligos 204+206 mix (10µM each) | 1 |
| DreamTAQ (5U/µL) | 0.2 |
| template (10ng/µL) | 2 |
| Nuclease free water | 14.3 |
| total | 20 |
| Cycling conditions | |

**95°C 3min 95°C 30sec, 58°C 20sec, 72°C 30sec (26 cycles) 72°C 5min**

**mutant *cmt3* allele**

| Component | volume (µL) |
|---|---|
| HF bufffer (5X) | 4 |
| dNTPs (10mM each) | 0.5 |
| Oligos 205+206 mix (10µM each) | 1 |
| Phusion (2U/µL) | 0.2 |
| template (10ng/µL) | 2 |
| Nuclease free water | 12.3 |
| total | 20 |
| Cycling conditions | |

**98°C 30sec 98°C 10sec, 65°C 30sec, 72°C 1min (26 cycles) 72°C 5min**

**WT *KYP* allele**

| Component | volume (µL) |
|---|---|
| Green buffer (10X) | 2 |
| dNTPs (10mM each) | 0.5 |
| Oligos 201+203 mix (10µM each) | 1 |
| DreamTAQ (5U/µL) | 0.2 |
| template (10ng/µL) | 2 |
| Nuclease free water | 14.3 |
| total | 20 |
| Cycling conditions | |

95°C 3min 95°C 30sec, 58°C 20sec, 72°C 30sec (25 cycles) 72°C 5min

**mutant *kyp* allele**

| Component | volume (µL) |
|---|---|
| Green buffer (10X) | 2 |
| dNTPs (10mM each) | 0.5 |
| Oligos 202+203 mix (10µM each) | 1 |
| DreamTAQ (5U/µL) | 0.2 |
| template (10ng/µL) | 2 |
| Nuclease free water | 14.3 |
| total | 20 |
| Cycling conditions | |

95°C 3min 95°C 30sec, 58°C 20sec, 72°C 30sec (25 cycles) 72°C 5min

***NRPE1* allele**

| Component | volume (µL) |
|---|---|
| HF bufffer (5X) | 4 |
| dNTPs (10mM each) | 0.5 |
| Oligos 275+276 mix (10µM each) | 1 |
| Phusion (2U/µL) | 0.2 |
| template (10ng/µL) | 2 |
| Nuclease free water | 12.3 |
| total | 20 |
| Cycling conditions | |

98°C 30sec 98°C 10sec, 60°C 20sec, 72°C 30sec (35 cycles) 72°C 5min

**Table S4.** PCR conditions used for all genotyping carried out in this study.

| Fragment | Name | Primer Sequence[a] | Size (bp)[b] | Tm°C |
|---|---|---|---|---|
| **Rj** | KL1310 | TGGTGTTTAAAAATTYATGTTTTTGTG | 320 | 50 |
| | KL1844 | TCCACRARTCATCRTCTCCAAACA | | |
| **Fie2** | KL1426 | AAGATTTGAGATTYGATTTGAAGTGTG | 225 | 52 |
| | KL1427 | CTTTCCCCTCCRCCTAATTCTCCTTA | | |

**Table S5.** Oligonucleotides used for maize experiments.

| Sample | Total reads sequenced | Total mapped reads | Total mapped reads after removing PCR duplicates | Total Hi-C reads after filtering |
|---|---|---|---|---|
| **Replicate 1** | | | | |
| **M82** | 256,921,384 | 149,308,782 | 121,182,766 | 47,391,805 |
| ***sulf*** | 197,852,756 | 116,087,535 | 96,693,269 | 34,996,574 |
| **Replicate 2** | | | | |
| **M82** | 56,174,862 | 32,892,122 | 32,181,207 | 13,128,114 |
| ***sulf*** | 55,212,666 | 31,349,383 | 30,419,079 | 12,043,402 |
| **Replicate 3** | | | | |
| **M82** | 64,995,720 | 39,275,083 | 38,441,719 | 14,651,575 |
| ***sulf*** | 37,915,680 | 23,337,027 | 22,656,587 | 9,378,235 |

**Table S6.** Number of Hi-C reads (statistics on mapping, PCR duplicates, and finally retained true Hi-C reads).
